# Engineering sialylation in pigs: A promising strategy for overcoming xenograft rejection and revolutionizing organ transplantation

**DOI:** 10.1101/2025.04.30.651299

**Authors:** Saptarshi Roy, Chaitra Rao, Emily K. Sims, Rita Bottino, David K.C. Cooper, Jon D. Piganelli

## Abstract

The organ shortage crisis leaves over 100,000 people waiting for transplants, causing 6,000 deaths annually. To address this, pigs are being explored as potential donors. Despite advances like the FDA-approved GalSafe pig, immunological challenges remain. Key issues include strong antibody, innate, and cellular immune responses, along with coagulation problems due to differences in glycosides and sialic acid linkages, which prevent long-term xenograft survival. Hyperacute rejection, caused by instant blood-mediated immune reaction (IBMIR), is a persistent problem, characterized by inflammatory and thrombotic responses when, for example, xenogeneic islets contact blood or reperfusion post anastomosis of transplanted organ. To overcome IBMIR, and other innate immune mediated rejection, we expressed human sialyltransferase (ST8Sia6) in otherwise wild type, porcine endothelium, creating human-like sialic linkages on porcine glycoproteins and lipids. This increased expression of human sialic acid 2,8- linkage on porcine endothelial cells, enhanced sialic-acid-binding immunoglobulin-like lectin (Siglec) binding reducing immune effector mechanisms such as complement activation and cell cytotoxicity. It also correlated with reduced CTL-targeted killing, lower levels of CD107a, perforin, and IFN-γ production. This coincided with higher immunoreceptor tyrosine-based inhibition motif (ITIM) induction, mirroring immune tolerance seen in fetal development and tumor immune evasion. Moreover, the landscape of the expression induced transcriptome of ST8Sia6, overexpression in porcine kidney cells revealed differential expression of genes involved in immune downregulation, cell signaling, and metabolic alteration. Expression of α2,8-linked disialic acids on porcine cells protected against immune effector mechanisms, reducing complement activation, immune cell activation, and CTL killing. These findings suggest that enhanced α2,8- linked disialic acid expression can modulate the innate and adaptive immune response, reducing xenograft rejection. This approach may improve xenotransplantation success, mitigate primary non-function in xenografts, and be applied to human iPSC-derived islets and other cell products. Further research into the specific mechanisms of these immunomodulatory effects could guide the development of effective strategies for xenotransplantation.

## Introduction

The disparity between the demand for organ transplants and the availability of suitable human donors remains a significant challenge in healthcare. Recent data from Health Resources & Services Administration, over 103,000 individuals are on the national transplant waiting list in the United States ^1^. Unfortunately, 6000 people a year die waiting for an organ transplant. To overcome transplantation reliance on the availability of a limited number of human donors, which is a major constraint, xenogeneic species such as pigs are being considered as potential tissue donors ^2^. According to the Organ Procurement and Transplantation Network (OPTN), in 2023, a record 46,632 transplants were performed, yet the number of patients awaiting transplants continues to exceed the supply. Several factors contributing to the imbalance between supply and demand is the escalating need for organ transplants. The prevalence of end-stage renal disease in the United States has surged, driven by a sharp rise in hypertension, obesity, and diabetes within the population ^3^. Numerous patients are on waiting lists because of the demand for organ transplants continuing to far outpace the supply of qualified human donors, underscoring the urgent need for alternative approaches. One of the most important developments in contemporary medicine is organ transplantation, which offers patients with end-stage organ failure life-saving options ^4^.

To overcome transplantation reliance on the availability of a limited number of human donors. A promising solution lies in xenotransplantation, the practice of transplanting organs or tissues from pig to human ^5^. This approach offers the potential to significantly expand the pool of available donor organs, addressing the shortage and providing hope to patients in need of life-saving procedures ^6^. However, significant barriers still exist for successful xenotransplantation. The immunological rejection caused by molecular incompatibilities between human and porcine cells^7^. In an effort circumvent these incompatibilities, the U.S. Food and Drug Administration approved a first-of-its-kind intentional genomic alteration (IGA) in a line of domestic pigs, referred to as GalSafe pigs, which are disrupted in the glycoprotein galactosyltransferase alpha-1,3 (GGTA1) gene. This gene normally codes for an enzyme that is responsible for production of the alpha-gal sugar (galactose-alpha-1,3-galactose) found on biological surfaces, such as cells, tissues, and organs in all mammals except humans and certain nonhuman primates, alpha-gal on the donor tissue can trigger a strong immune rejection. In homozygous GalSafe® pigs, the GGTA1 gene is knocked out on both alleles, resulting in the intended trait of no detectable alpha-gal sugar on their cells, tissues, or organs making these organs more compatible for human transplants ^8^. Pigs containing the disrupted GGTA1 gene pass the trait to their offspring through conventional animal breeding.

From these first modifications advances came rapidly with genetically engineered pigs, starting with the introduction of a human complement-regulatory protein (CRP) gene ^9^. Decades of progress in xenotransplantation, propelled by the advent of rapid genome editing techniques, most notably CRISPR-Cas9 technology, have ushered in extraordinary advancements ^10,11^. These breakthroughs in large animal preclinical models laid the foundation for clinical trials in the United States. In 2003-2004, combination galactosyltransferase gene-knockout and expression of a human CRP protect early antibody mediated organ rejection. Using this highly advanced technology, “triple knock-out” (TKO) pigs, which lack three major carbohydrate xenoantigens (αGal, Neu5GC, and SDa), have been recently created ^12^. These pigs may be used for food or human therapeutics, ensuring the production, regulation, safety, and control of this form of livestock is well prepared to meet this organ donation need. Even with recent technological advances, such as the GalSaf pig, several immunological hurdles remain ^13^. For example, xenotransplantation faces challenges due to strong antibody-mediated immune responses and coagulation issues, which hinder long-term graft survival. Antibody-driven complement activation triggers inflammation, endothelial damage, and coagulation cascade activation, even under the full accompaniment of front-line immunosuppression ^14–17^. This leads to interstitial hemorrhage, thrombus formation, ischemia, and eventual graft failure ^18,19^. Specifically, hyper acute rejection resulting from instant blood mediated immune reaction (IBMIR) continues to plague the field. IBMIR is characterized by a nonspecific inflammatory and thrombotic reaction where xenografts encounter blood, even in GalSaf modified pigs ^20^.

To overcome the instant blood-mediated inflammatory reaction (IBMIR) and the innate-mediated targeted tissue destruction, we describe a novel approach involving the expression of human Sialic acid (a nine-carbon monosaccharides) in porcine endothelium surface. The Sialyltransferases are enzymes that transfer sialic acid to nascent oligosaccharides of glycoprotein as a transmembrane protein, four major types of glycosidic bonds can be formed, namely, Neu5Acα2-6Gal, Neu5Acα2-3Gal, Neu5Acα2-6GalNAc, and Neu5Acα2-8Neu5Ac ^21^. Neu5Ac and Neu5Gc are the predominant forms in mammals. Since humans are unable to convert Neu5Ac into Neu5Gc due to a mutation in the cytidine monophosphate-N-acetylneuraminic acid hydroxylase gene, only Neu5Ac is produced de novo ^22^. Each sialyltransferase is specific for a particular sugar substrate and serves to transduce signals via interaction with sialic-acid-binding immunoglobulin-like lectins (Siglecs) on immune cells specifically NK cells, T cells and macrophages ^23^. This sialic acid-Siglec interaction has a profound impact on the ability of innate immune cells to become activated by evoking an inhibitory signal to surveilling, natural killer (NK), CD4, CD8 T cells, and macrophages, as well as inhibiting the binding of natural IgM antibodies and complement to suppress rapid xenograft rejection by adding sialic acid residues to the cell surface, mimicking human cell markers. By expressing, human sialic acid moieties on the surface of porcine endothelia we can circumvent the activation of innate immune cells and the binding of natural antibodies and complement to porcine endothelia, essentially cloaking these cells from immune recognition ^24^. Interestingly, this type of immune modulation mirrors mechanisms observed in both fetal tolerance during pregnancy and within the tumor microenvironment. In these contexts, immune evasion is facilitated through the expression of self-like glycan structures and the creation of an immunosuppressive milieu, allowing the fetus or tumor to persist despite the host’s immune surveillance ^25,26^. Similarly, engineering porcine endothelia to display human-like sialylation patterns mimics this natural strategy of immune invisibility, offering a promising avenue for reducing xenograft rejection ^27,28^.

There are six sialyltransferases, preferentially used by humans, ST8Sia1 through ST8Sia6. Among them, ST8Sia6 is distinct in its ability to synthesize disialic acids by attaching a single α2,8-linked sialic acid to an existing α2,3- or α2,6-linked sialic acid. In contrast, ST8Sia1 through ST8Sia5 primarily facilitate the addition of α2,8-linked sialic acids to glycoproteins and glycolipids, playing crucial roles in neural development, immune responses, and cell-cell communication ^29,30^. The unique function of ST8Sia6 in generating disialic acids significantly influences protein structure and function. These disialic acids have been linked to various diseases, including cancers and neurological disorders, underscoring the importance of studying the regulation and role of this specialized enzyme in immune inhibition ^31^. Previous research by scientists has shown that the sialyltransferase ST8Sia6 produces ligands for Siglecs expressed on immune cells ^32^. These ligands inhibit inflammation by recruiting protein tyrosine phosphatases to the immunoreceptor tyrosine-based inhibition motif (ITIM), which is subsequently followed by the recruitment of distinct SH2-containing molecules to the receptor site. These proteins negatively regulate receptor-linked signal transduction pathways ^24,33^. To study immune evasion, we overexpressed the sialyltransferase ST8Sia6 to attach α2,8-linked sialic acid to existing α2,3- or α2,6-linked sialic acids on the porcine cell membrane. This essentially obscures the α2,3- or α2,6- linked sialic acids on the surface glycoproteins of porcine kidney and aortic endothelial cells, which human immune cells recognize as foreign. We hypothesized that this strategy would inhibit human immune cells, blocking recognition and activation of the immune system against the xenograft tissue, thereby prolonging xenograft survival by evading the initial immune response triggered by IBMIR. The strategy was designed to prolong xenograft survival by creating a more hospitable environment that evades the initial immune response triggered by the IBMIR. This sialic acid modifications mark cells as “self” by adding sugar moieties recognized by the immune system, preventing complement-mediated destruction during xenotransplantation. This mechanism normally maintains tolerance, to self-recognition, but also causes graft rejection when foreign tissues, like porcine organs, lack these markers. Similarly, recent thymic emigrants (RTE) rely on proper-sialylated sugars to signal self-recognition via Siglec receptors. Defective sialylation in RTE results in immune targeting and destruction, by an antibody-complement mediated mechanism, as they are perceived as non-self ^34^. Furthermore, sialic acid-Siglec interaction inhibits targeted immune responses, thwarting rejection of the fetus during pregnancy and in tumors. Increased glycolysis in cancer cells boosts sialic acid precursors, enhancing glycoprotein sialylation and immunosuppression of tumor-infiltrating lymphocytes. Hypersialylation of 40-60% of tumor cell surfaces aids immune evasion and tumor survival. In the placenta, sialic acid modulates immune tolerance, preventing maternal immune attacks and supporting trophoblast invasion, cell communication, and placental development for pregnancy success ^25^. Recently others have taken the opposite approach with incredible success by expressing the recombinant porcine a1,3GT gene in Newcastle disease virus (NDV-GT) to essentially induce the IBMIR triggered hyperacute rejection of tumors expressing the NDV-GT ^35^ .

Thus, exploiting this strategy to express sialyltransferases in porcine endothelia cells to cloak the porcine tissue from the innate immune response may have a protective benefit that has never been explored in the current literature. To examine this hypothesis, we assessed the effects of expressing the sialyltransferase ST8Sia6, on porcine kidney and aorta endothelia, which adds an α2,8-linked sialic acid linkage on the surface of the porcine cells. We then assessed this interaction with immune cells, in different immune mediated cytotoxic functions, immune cytokine pathway, regulation of complement activation, involvement of ITAM and ITIM, and transcriptomic alterations. This approach should provide insights into the potential mechanisms by which sialyltransferase expression could modulate the innate immune response and promote xenograft tolerance. The comprehensive transcriptomics study serves as a map to help guide the impact and limitations of ST8Sia6 gene expression in regulating the xeno-immune response. These targeted changes will inform us and guide the development of adjunct therapies to address shortcomings in the treatment to help circumvent acute hyper rejection, achieve durable graft tolerance, and manage chronic rejection. Understanding these interactions, and the changes in the transcriptomic landscape will educate us in further refining the development of novel strategies to enhance xenograft survival.

## Results

### Overexpression of ST8Sia6 in mice melanoma cell lines precipitate ligands for Siglec-E

In many cancer types, hypersialylation marked by increased incorporation of sialic acids into tumor cell surface glycans is a common feature aimed at evading immune surveillance ^36,37^. These sialic acids can interact with sialic acid immunoglobulin-like lectins (Siglecs) and serve to assess self-non-self-tissues during immune surveillance ^28,38^. They are added to glycans through one of three linkages, named after the carbon atom on the terminal sugar to which the sialic acid attaches (α2,3, α2,6, or α2,8), catalyzed by one of twenty different sialic acid transferases ^29,39–41^. Public databases show that ST8Sia6 is upregulated in human cancers, correlating with changes in cellular proliferation and viability ^42^. The well-characterized B16-F10 and B16-Ova melanoma cells do not exhibit detectable levels of ST8Sia6 ^42^. To investigate its role in immune modulation, we engineered paired cell lines with and without ST8Sia6 expression. We then analyzed the expression of ligands for murine Siglec-E during stable ST8Sia6 expression. GFP-tagged ST8Sia6 plasmids were transfected into B16-F10 and B16-Ova cells, and ST8Sia6 expression was detected by observing GFP expression through a fluorescent microscope at different time points (Supplementary Fig. 1a). As a control the same parent plasmid was used carrying only the reporter GFP plasmid. This was transfected into the cell lines as controls. Highest expression of ST8Sia6 was found in B16-F10 and B16-Ova cell lines within 24hr of post transfection (Fig. 1a). We also estimated the frequency and transfection efficiency estimated by flowcytometry (Supplementary Fig. 1d-e), which represented the percentage of GFP transfected cells (Fig. 1b). Furthermore, Western Blot analysis demonstrated a significant overexpression of ST8Sia6 protein from B16-F10 and B16-Ova transfected cell lysate compared to reporter control (Fig. 1c). To assess the functional impact of ST8Sia6 expression, recombinant fluorescently-tagged-Siglec-E, which binds to α2,8-linked disialic acids was used to assess both expression and binding. We employed this system to detect ligand expression on B16-F10 and B16-Ova cells. In this study we demonstrated that, B16-F10 and B16-Ova cell lines transfected with ST8Sia6 exhibited significantly enhanced binding frequency with recombinant Siglec-E compared to cells transfected with control GFP plasmid (Fig. 1e-f). Notably, Siglec-E also exhibited binding to the control GFP- transfected cell lines, albeit to a lesser extent, indicating recognition of sialic acid products generated by other sialyltransferases ^43,44^. Lectin binding assays using peanut agglutinin (PNA) were conducted to further explore the impact of ST8Sia6 expression. The results revealed that B16-F10 and B16-Ova cell lines transfected with ST8Sia6 exhibited significantly reduced PNA binding compared to control GFP-transfected cells (Fig. 1d, Supplementary Fig. 1b). This reduction suggests that ST8Sia6-mediated addition of α2,8-linked sialic acid interferes with the recognition of O-linked oligosaccharides by PNA, indicative of cloaking other sialic acid linkages from immune surveillance. Conversely, control GFP-transfected cells showed preferential binding to PNA, indicating the presence of accessible O-linked oligosaccharides for PNA recognition. These findings underscore the role ST8Sia6 expression plays in immune modulation and its potential as a therapeutic target in various immune regulatory treatment strategies.

**Figure 1.**
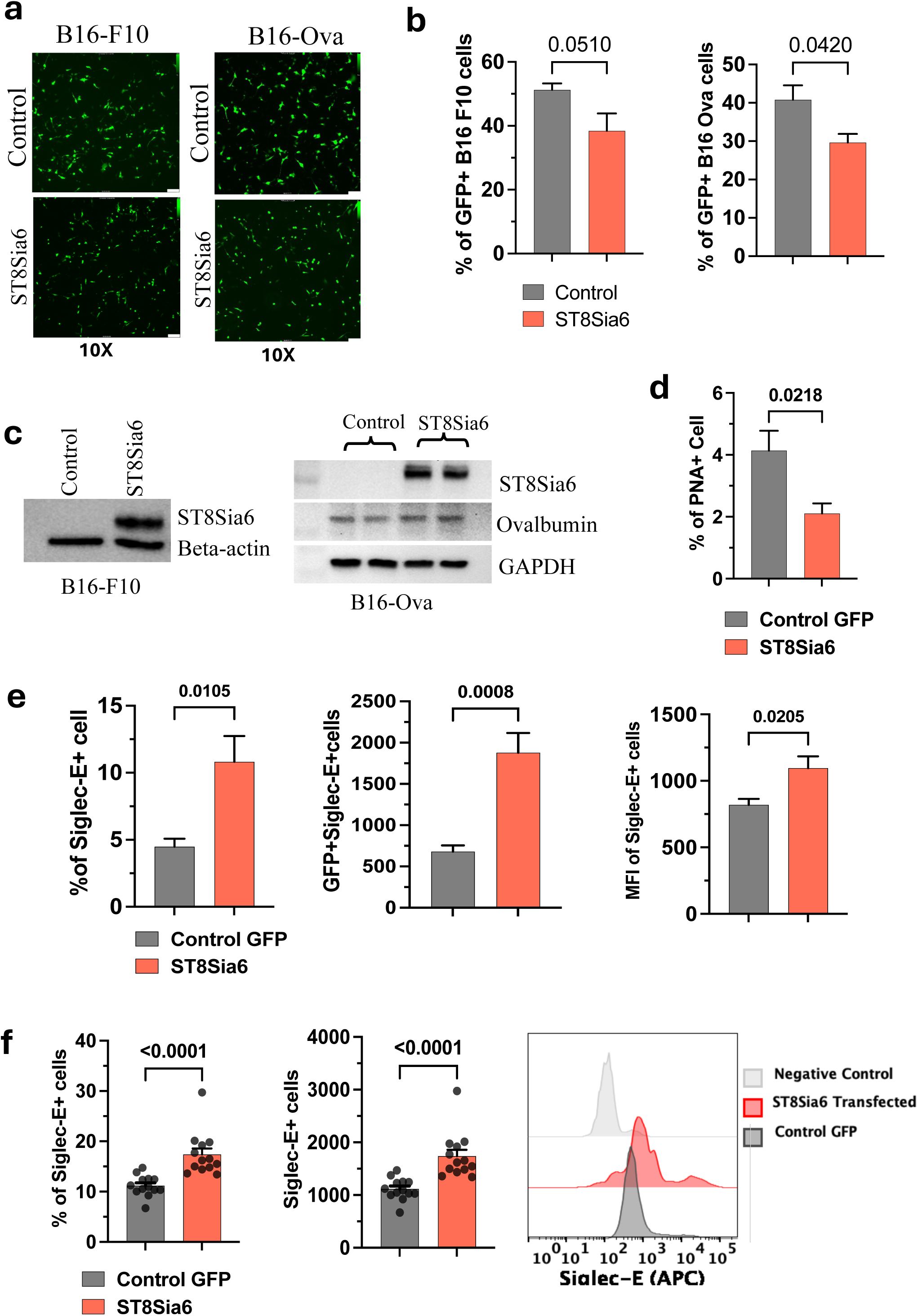
Overexpression of ST8Sia6 in B16-F10 and B16-Ova cell lines, and estimating ligands generation for Siglec-E binding. a, GFP fluorescent images showed transfection efficiency of ST8Sia6 in B16-F10 and B16-Ova cell line. **b,** Assessment of transfection efficiency using flow cytometry in B16-F10 cells. **c,** Lysates from B16-F10 and B16-Ova transfected cells were examined for expression of ST8Sia6 by Western blotting. The molecular weight of ST8Sia6 is 43 kDa. **d**, Peanut Agglutinin (PNA) lectin binding assay in ST8Sia6 plasmid or Control GFP transfect B16-F10 cell line. **e-f,** B16-F10 and B16-Ova murine cancer cell lines were stably transfected with ST8Sia6 plasmid or Control GFP and were probed for ligands with recombinant Siglec-E, expression was measured by flowcytometry using anti-Siglec-E antibody. Siglec-E ligand positive cells percentage, numbers and geometric mean fluorescence intensity (MFI) was quantified across three (n-3) independent experiments, and significance analyzed by two-tailed unpaired t test between groups, significant different (P<0.05).

### Stable expression of ST8Sia6 in melanoma cells correlates with immune suppression and reduces T cell-driven cytotoxicity

To determine the effect of ST8Sia6 expression on immune modulation, we performed an in vitro cytotoxic killing assay, using the murine, ova-peptide specific, OT-1 CD8+ T cell. OT-1 T cells were co-cultured with ST8Sia6 transfected B16-F10 pulsed with K^b^-specific class I ova-peptide SIINFEKL and ST8Sia6 plasmid transfected B16-Ova cells individually at target (B16-F10/ B16- Ova) and different effector (OT-1 CD8+ T) cell ratio for 24 h (Fig. 2a). After 24 hr. coculture, we analyzed the killing efficiency of OT-1 CD8+ T cells on ST8Sia6 transfected B16-F10 and B16- Ova cells. Our results revealed a significant differences in relative killing efficiencies, correlating with a difference in the relative number of live B16-F10 cells were observed between the control GFP and ST8Sia6 plasmid-transfected groups. OT-1 CD8+ T cells showed in average 60% killing on ST8Sia6 transfected B16-F10 cell (pulsed with SIINFEKL) and B16-Ova cells respectively, whereas control group showed more than 60% killing in both cell lines (Fig. 2b, Supplementary Fig. 2a). Further flowcytometry analysis revealed that a significantly higher number of GFP+ cells were present in ST8Sia6 transfected group in comparison with control (Fig. 2c). To assess the full cytotoxic potential of OT-1 CD8+ T cells, we determined the effector-dependent cytokine IFN-γ expression level from experimental culture supernatant. Expression of ST8Sia6 transfected B16- F10 tumor cells elicited a reduced expression of IFN-γ OT-1 CD8+ T cells compared to control GFP group (Fig. 2d). Similarly, we found a significant reduction in IFN-γ production in B16-Ova ST8Sia6 transfected cells during co-culture condition compared to control (Supplementary Fig. 2b). To investigate the expression of OT-1 CD8+ T cell effector subpopulations, we collect the OT-1 CD8 T cells from different groups of cocultured cells and performed flowcytometry stanning for early activation and cytotoxic effector molecules, which synergize to mediate apoptosis of target cell. CD69 and LAMP-1 activation marker expression by Flow cytometry analysis showed that ST8Sia6 transfection in B16-F10 and B16-Ova cells led to a significant reduction in the expansion of CD8+CD69+ and CD8+LAMP-1+ T cell subsets compared to control-GFP transfected cells (Fig. 2e-j). Specifically, the absolute number of CD8+CD69+ and CD8+LAMP- 1+ cells was significantly lower in B16-F10 (Fig. 2f and 2i) and B16-Ova (Supplementary Fig. 2f) cells transfected with ST8Sia6 were used as targets. Additionally, the median fluorescence intensity (MFI) of CD8+ T cell activation markers CD69 and LAMP-1 respectively were significantly lower in the ST8Sia6-transfected B16-F10 cell co-culture conditions (Fig. 2g and 2j), similarly MFI of CD8+CD69+ cells and CD8+LAMP1+ cells significantly low in ST8Sia6 transfected B16-Ova experiment condition (Supplementary Fig. 2e). Results from various E:T ratio model revel that, concentration dependent increase of killing efficiency of OT-1 T cells was observed in B16-F10 and B16-Ova control GFP and ST8Sia6 plasmid transfected cells. Different target effector conditions in B16-F10 ST8Sia6 transfected cells also revealed a gradual increase of IFN- γ secretion in presence of OT-1 CD8. Collectively, these results demonstrate that ST8Sia6 overexpression contributes to inhibiting the overall immune activation and effector CTL killing function against the target cell, thus providing potent immunosuppression against the target tissue, making it a potential target for immunotherapy aimed at maintaining immune homeostasis in the microenvironment.

**Figure 2.**
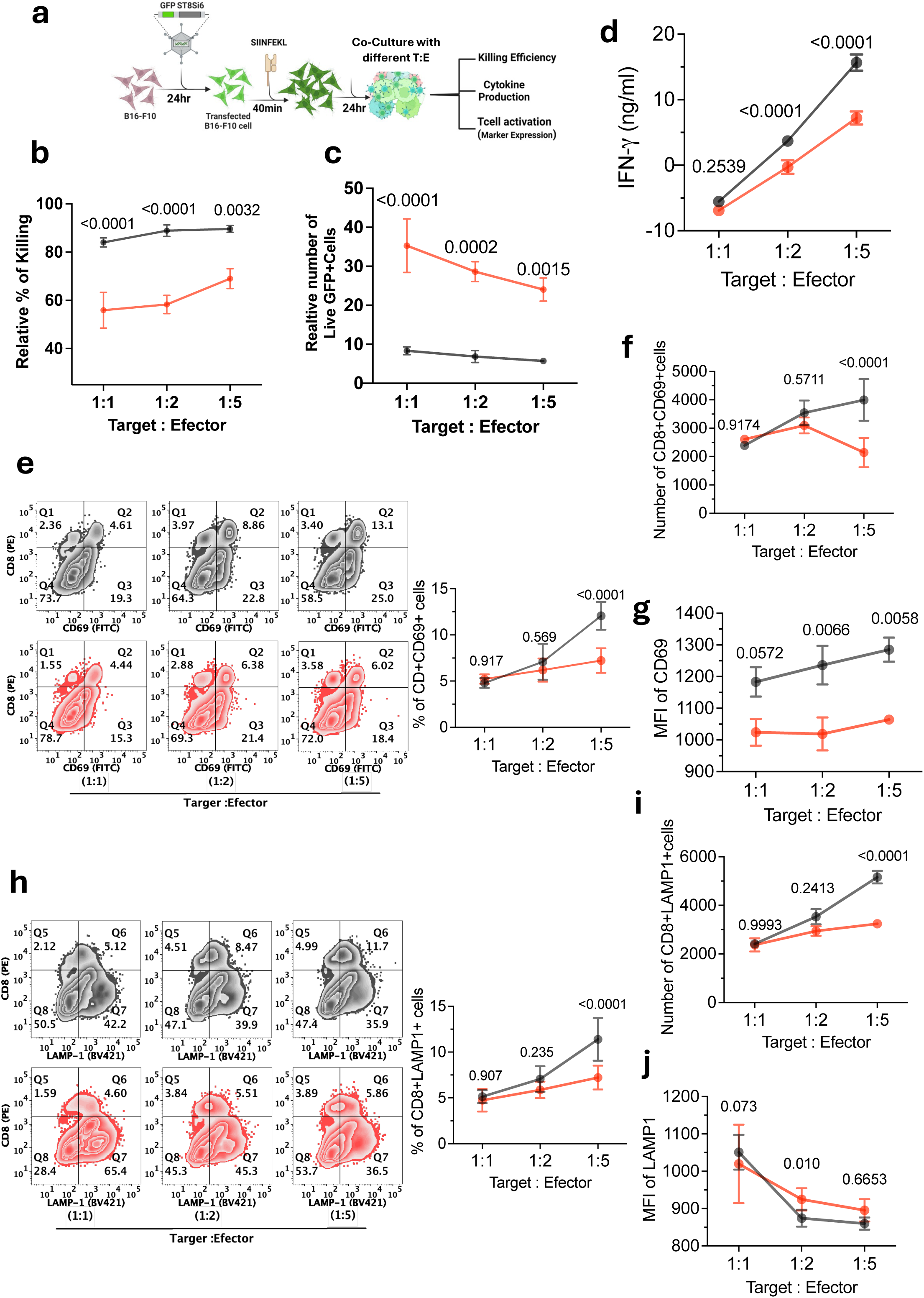
**Overexpression of ST8Sia6 in B16-F10 and B16-Ova cells suppress OT-1 CD8+ T cells mediated cytotoxic effects**. In this experimental conditions, B16-F10 cells transfected with a control GFP plasmid (grey curve) and those transfected with ST8Sia6 plasmid (red curve) were cocultured with OT-1 CD8+ T cells, which are specific effector cells, at various effector to target (E:T) ratios. The cytotoxicity was assessed relative to maximal cytotoxic effects. **a,** Schematic representation of in vitro cytotoxic killing assay, where OT-1 CD8+ T cells were co-cultured with ST8Sia6 in B16-F10 pulse with SINFEKL and B16-Ova cells in individual setup at a T cell: B16 cell at different ratio for 24 h. **b,** The frequency of killing efficiency, and c, relative number of viable GFP-expressing target cells indicate the extent of effector cell-mediated cytotoxicity across different (E:T) ratios. **d,** IFN-γ levels were evaluated using ELISA from the supernatant of cocultures at different (E:T) ratios. **e and h,** Flowcytometry counter plot represent the percentage of CD8+CD69+ cells and CD8+LAMP1+ cells population at various effector to target (E:T) ratios conditions. **f,** Number of CD8+CD69+ cells. **g,** MFI of CD69 expression in CD8+T cells across different E:T conditions. **i,** Number of CD8+LAMP1+ cells. **j.** MFI of LAMP1 expression in CD8+T cells across different E:T conditions. Data is representative of three independent experiments (mean ± SEM). E: T ratio = effector: target ratio; MFI = Mean Fluorescence Intensity. Significant difference assessed by two-way ANOVA and multiple comparison test between groups, significant different (P<0.05).

### Transfection of porcine kidney and aorta endothelial cells with ST8Sia6, to assess expression of 2,8-linkage generation and binding to specific receptors, Siglec-E, Siglec-9, and Siglec-10

To evaluate the impact of 2,8-linked sialic acid expression on porcine cell interactions with human peripheral blood cells, we assess the interaction with selected Siglec receptors, as described above by flow cytometry and then using human immune cells, isolated from peripheral blood [23]. These studies aim to dissect the immunomodulatory role of 2,8-linked sialic acid expression on porcine cells and its involvement in Siglec-mediated inhibitory signaling. The findings could have significant implications for understanding xenograft rejection and reducing immune responses in xenotransplantation. As described above for the B16 model of targeted expression and recognition, porcine kidney and aorta endothelial cells were transfected with the ST8Sia6 gene, which included a green fluorescent protein (GFP) reporter, and a control plasmid expressing only GFP (Fig. 3a and f). Transfection efficiency was assessed using flow cytometry, showing 55% for porcine kidney cells and 50% for porcine aorta endothelial cells (Fig. 3c, h). Additionally, substantial ST8Sia6 protein expression was detected in cell lysates from the transfected porcine kidney and aorta cells (Fig. 3b, g). To examine the functional impact of ST8Sia6 expression, fluorescently labeled-recombinant Siglec-E, Siglec-9, and Siglec-10 were used to assess ligand binding on the transfected cells. As expected, Siglec-E (Fig. 3d, i) and Siglec-9 (Supplementary Fig. 4a, b) showed significantly higher binding to ST8Sia6-transfected porcine kidney and aorta cells compared to control-GFP conditions. The number of GFP+ cells binding to Siglec-E and Siglec-9 was also significantly higher in ST8Sia6-transfected cells (Fig. 3e, j and Supplementary Fig. 4c, d). However, no significant differences were observed in Siglec-10 binding in porcine kidney and aorta endothelial cells, confirming the specificity of the ST8Sia6-dependent receptor-ligand interaction (Supplementary Fig. 4a-d). These findings demonstrate that forced expression of ST8Sia6 promotes specific 2,8-linkage, facilitating binding with Siglec-E and Siglec-9 receptors. This 2,8-linkage expression on porcine endothelial cells is likely to influence the immune inhibitory effects when these cells are cultured with human peripheral blood lymphocytes. The interaction between porcine endothelial cells and human lymphocytes could impact xenotransplantation by modulating immune responses through the 2,8-linkage, potentially reducing rejection risk. Further research is needed to explore the mechanisms and clinical applications.

**Figure 3.**
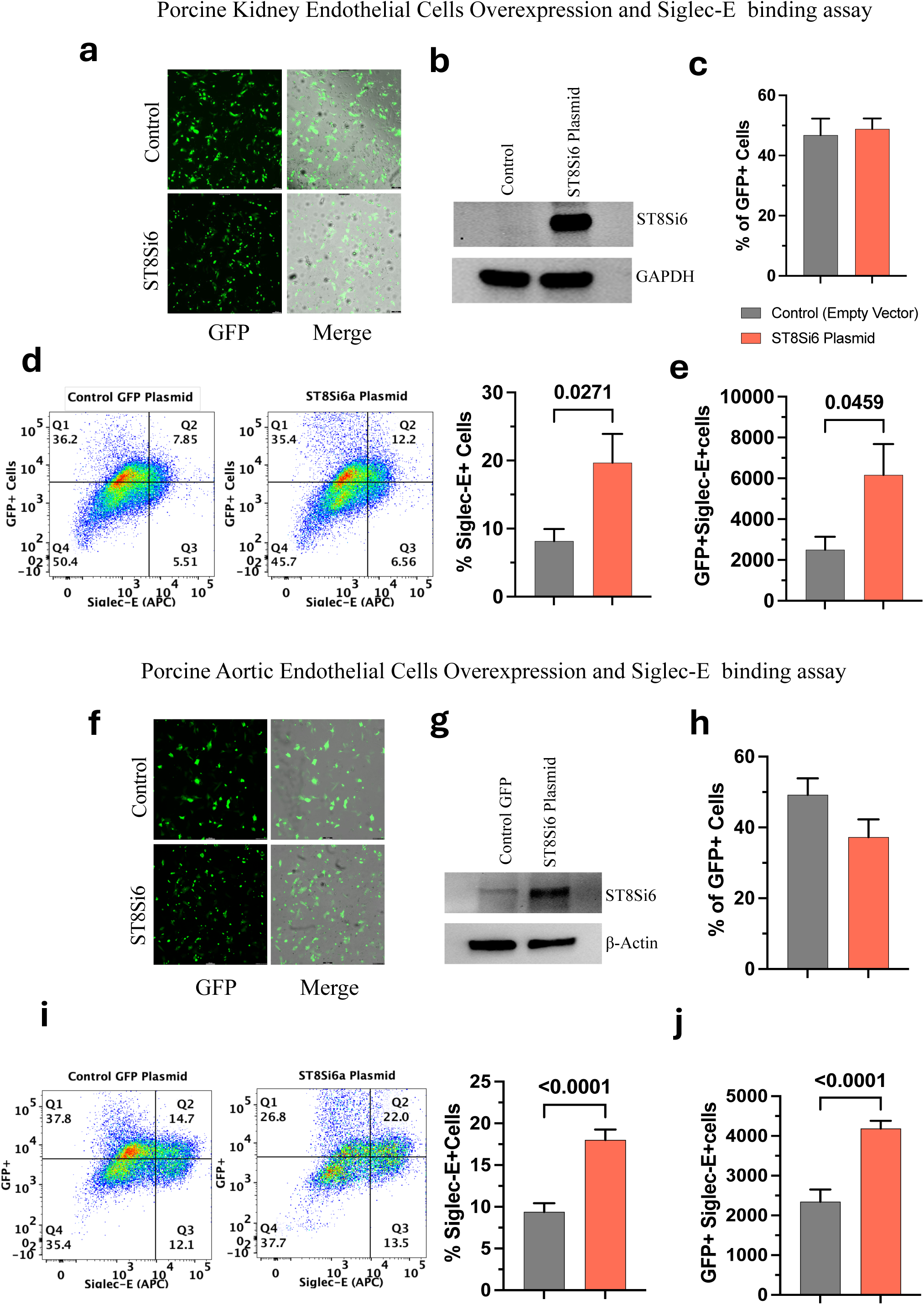
Effect of ST8Sia6 overexpression on Siglec-E ligand expression in primary porcine kidney and aortic cell lines. a & f, GFP fluorescent images demonstrated the transfection efficiency of ST8Sia6 in primary porcine kidney and aorta endothelial cell lines. **b & g**, Western blot analysis indicated that overexpression of the ST8Sia6 gene increased ST8Sia6 protein levels in primary porcine kidney and aorta cell lines, but no expression in control GFP plasmid transfected cells. **c & h**, Evaluation of transfection efficiency using flow cytometry in porcine kidney and aorta endothelial cells. Siglec-E binding assays were performed by probing for ligands using recombinant Siglec-E, and the expression levels were assessed via flow cytometry with an anti-mouse Siglec-E antibody. **d & I,** Percentage of Siglec-E ligands positive porcine kidney and aorta cells was represented in scatter plots and bar graphs. **e & j,** Additionally, the absolute number of Siglec-E positive porcine kidney and aorta endothelial cells was quantified using flow cytometry. All data are represented in four independent experiments and presented as mean ± SE, significance analyzed by two-tailed unpaired t test between groups, significant different in P value (P<0.05).

### Immune cell cytotoxic effects are blunted in ST8Sia6 expressing porcine kidney and aorta endothelial cells

Porcine kidney and aorta endothelial cells expressing the sialyltransferases ST8Sia6 (2,8-linkage sialic acid) were exposed to healthy human PBMC cells. To evaluate xenograft reactivity, we studied human PBMCs exhibited cytotoxic effects, leading to increased viability of the sialic acid expressing porcine endothelial cells. This suggests that ST8Sia6 (specific 2,8-linkage) expression may render porcine cells more susceptible to protect immune-mediated rejection in xenotransplantation. Furthermore, to assess the potential of 2,8-linkage-Siglec-9 ligand expression to serve as a targeted therapeutic interaction for inducing immune cell inhibition, improving xenograft acceptance, we investigated whether ST8Sia6-expressing porcine kidney and aorta endothelial cells influence the downmodulation of immune cell-induced cytotoxic effects from human PBMCs. Therefore, we performed immune cell induce cytotoxic killing assay, where healthy human PBMCs were co-cultured with ST8Sia6 plasmid transfected porcine kidney and porcine aorta endothelial cells in individual setup at 1:5 target and effector cells ratio for 24 hr. (Fig. 4a). The results showed that ST8Sia6-expressing porcine endothelial kidney and aorta cells exhibited a statistically significant reduction in susceptibility to cytotoxic killing by human PBMCs compared to control-GFP transfected porcine kidney and aorta endothelial cells respectively (Fig 4.b and c). The maturation of CD4+ and CD8+ T cells to CTL effector function, is dependent on the expression of the proinflammatory cytokine Hu-IFN-γ, Hu-GrzB and Hu-perforin, secreted by cytotoxic T cells and natural killer cells. Therefore, we assessed the levels post killing assay. Consequently, to finds out the effects of ST8Sia6 expression in correlation with cytokine secretions, we measured Human IFN-γ, perforin and granzyme-B from culture supernatant. The results indicated a noteworthy decrease in the secretion of human IFN-γ, perforin, and granzyme-B by PBMCs when exposed to ST8Sia6-transfected porcine kidney and aorta endothelial cells respectively, in comparison to the control GFP transfected group (Fig. 4 d-i). These findings suggest that the expression of ST8Sia6 in porcine endothelial cells modulates the cytotoxic response of human immune cells by altering the secretion of key cytokine effector molecules. These results indicate that targeting the ST8Sia6-Siglec-9 axis has a strong impact on maturing CD8 T cells to CTL function which can be a potential strategy to mitigate immune-mediated rejection in xenotransplantation. Further investigation is warranted to fully elucidate the underlying mechanisms and explore the therapeutic potential of this approach.

**Figure 4.**
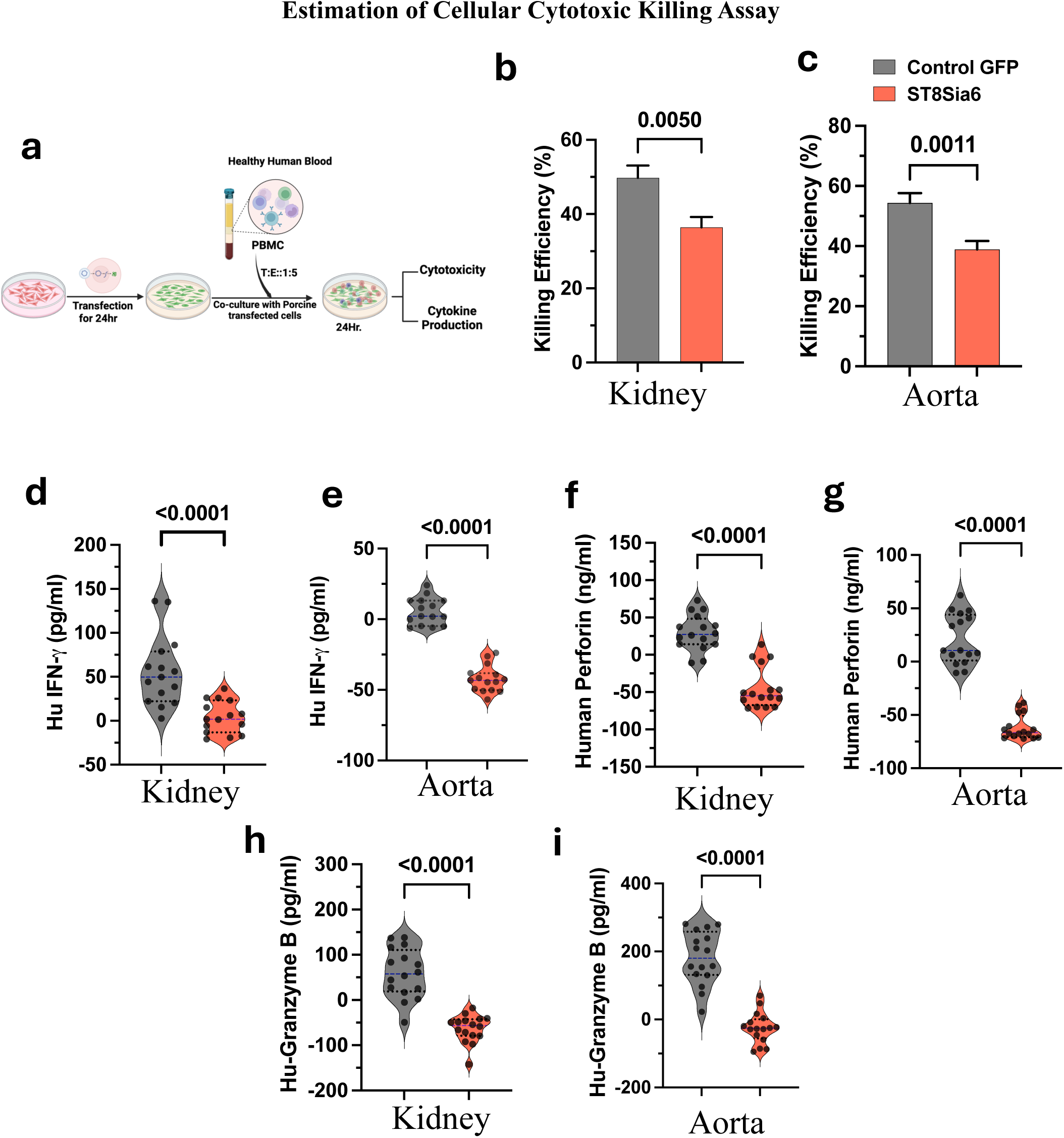
Overexpression of ST8Sia6 in porcine kidney and aorta endothelial cell line suppresses T and NK cells mediated immune cytotoxicity and cytokines production. a, Schematic illustration of the in-vitro experimental procedure, where ST8Sia6 transfected porcine kidney and aorta endothelial cells individually were co-cultured with healthy individual PBMC at 1:5 T:E ratio for 24 h (Target-Porcine cells, E-Human PBMC). Relative percentage of killing efficiency of PBMC against **b,** porcine kidney and **c,** porcine aorta endothelial compared with in control-GFP and ST8Sia6 overexpressed cells. T Cell activation was quantified by IFN-γ production by PBMC from **d,** porcine kidney, **e,** porcine aorta endothelial cells coculture supernatant. **f-g,** Hu-Perforin, and **h-i.** Hu-Granzyme-B cytokine productions by NK cells in PBMC are measured from supernatants after 24 hours of co-culture with porcine kidney and aorta endothelial cells respectively. All graphs are summarizing the result from 4 donors (mean ± S.E) with 4 individual replica. significance analyzed by two-tailed unpaired t test between groups, significant different in P value (P<0.05).

### Overexpression of ST8Sia6 in porcine endothelial cells mitigates the Instant Blood-Mediated Inflammatory Reaction in the presence of human blood

The Instant Blood-Mediated Inflammatory Reaction (IBMIR) is a major barrier to successful xenotransplantation, as it can lead to rapid rejection of the transplanted organ ^45,46^. The coagulation and complement cascades together with the leukocyte and platelet populations are the major players in IBMIR ^47^. This hyperacute innate immune attack dramatically affects primary xenograft failure and is driven by a robust innate immune response targeted by NK cells, macrophages and natural antibody, that can fix complement, leading to complement mediated cell death ^48^. By mitigating this reaction, ST8Sia6 overexpression could improve the viability and function of xenografts, potentially enhancing primary engraftment, thus increasing the prospects for sustained xenograft durability ^25^. To validate the effect of ST8Sia6 overexpression in IBMIR during xenotransplantation, we incubated ST8Sia6-transfected porcine kidney and aorta endothelial cells with heparinized healthy human whole blood for up to 6 hours, and blood was collected at different time point to determine the complement activation difference (Fig. 5a). Result indicates that, plasma C3a, C4b, C5b9 (membrane attack complex, MAC) complement factors significantly increase with time, when incubated with human blood in control transfected porcine kidney and aorta endothelial groups respectively. However, ST8Sia6-transfected porcine aorta endothelial cells effectively inhibit the elevation of these complement factors during incubation (Fig. 5c, e, g). In contrast, elevated plasma complement factors were observed in both ST8Sia6- transfected porcine kidney cells and the control groups group after incubation with human blood (Fig. 5b, d, h). Similar patterns were observed in plasma thrombin-antithrombin (TAT) complex, which is a key marker of coagulation activation and plays a vital role in regulating clotting. TAT complex formation also decreases in ST8Sia6-transfected porcine kidney and aorta groups compere to control group (Fig. 5h, and i). These findings suggest that overexpression of ST8Sia6 may reduce inflammatory responses associated with IBMIR, potentially improving xenotransplantation outcomes by inhibiting the activation of both complement and coagulation cascades. Overexpressing ST8Sia6 in porcine endothelial cells could help mitigate the IBMIR response, enhancing the xenografts survival and functions. Additionally, α2,8-sialyltransferase-6, in conjunction with Decay-Accelerating Factor (DAF), can prevent the formation of the membrane attack complex during complement activation. Similarly, CD46 (Membrane Cofactor Protein), a complement regulator, works with Sialyltransferase-8 to block the deposition of C3b, thus halting further progression of the complement cascade ^49,50^. Thus, further research is needed to fully understand how ST8Sia6 influences these pathways and to assess its potential in preclinical xenotransplantation models. Overcoming the IBMIR barrier is a crucial step toward realizing the potential of xenotransplantation as a viable solution for organ transplants.

**Figure 5.**
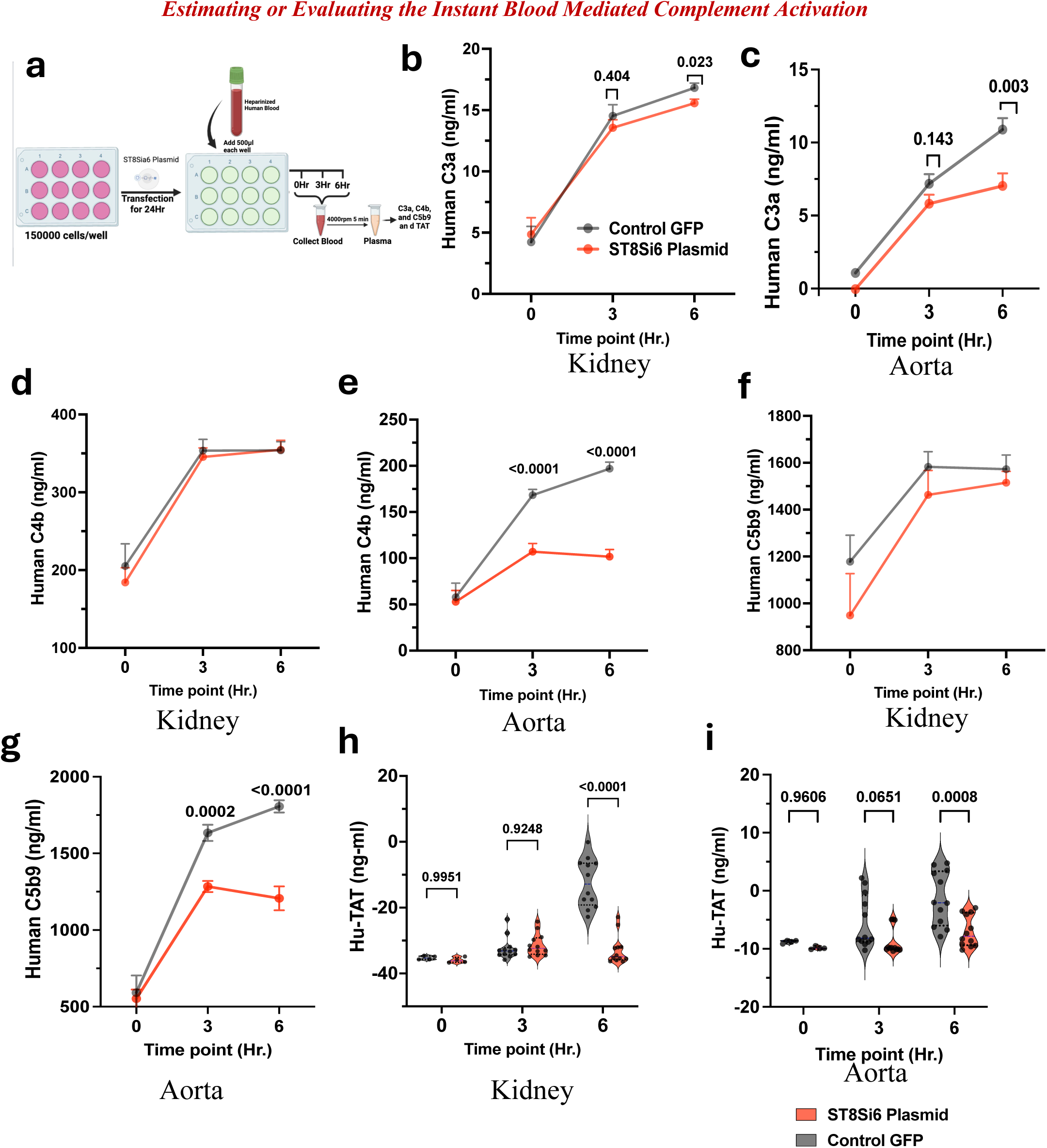
Evaluating valuated the instant blood-mediated complement activation. a, Schematic illustration of the in-vitro experimental procedure, measurements of human thrombin/anti-thrombin III complex (TAT), C3a, C4b, and C5b9 were taken from plasma in mixed human blood and porcine kidney and aorta tissues transfected with ST8Sia6. These were compared to controls porcine kidney and aorta endothelium transfected with GFP under in vitro conditions at three different time points (0, 3, and 6 hours). Result indicates that control GFP transfected Kidney and Aorta group (grey line) had increased complement activation. **b-c,** indicate C3a level in kidney and aorta condition respectively, similarly **d-e,** represent C4b level, **f-g,** represent C5b9 level from kidney and aorta experimental conditions. **h-i,** TAT level in plasma from kidney and aorta experimental conditions. In contrast, the ST8Sia6-transfected porcine kidney group (red line) showed a slight reduction in complement activation. Notably, the ST8Sia6- transfected aorta cells demonstrated significant reductions in complement activations. All graphs summarize the results from 4 donors (mean ± S.E) with 4 individual replicates. Significance differences in P value (P<0.05) were evaluated using ANOVA, and the P values are represented in the graphs.

### ST8Sia6 Overexpression in Porcine Endothelial Cells Modulates Immune Cell Activation and Surface Marker Expression on human PBMC Co-Culture Highlights NK and T Cell Responses

The phenotypic analysis of immune cells from ex vivo coculture experiments suggested that overexpression of ST8Sia6 regulates the functional activation and inhibition of T cells and NK cells ^50^. These effects influence T cell-mediated cytotoxic killing and the innate immune system, where natural killer (NK) cells play a critical role in the initial defense against xenograft rejections ^51,52^. Therefore, we evaluated the expression of several activation and inhibitory markers on lymphocytes from human PBMCs, in the presence of ST8Sia6-overexpressing porcine kidney and aorta endothelial cells, alongside control-transfected porcine cells (Fig. 6a). The results demonstrated a significant reduction in the frequency of T cells (CD3), NK cells (CD56), and natural killer T cells (CD3+CD56+) populations after 24 hours of coculture with ST8Sia6- overexpressing and control porcine kidney and aorta endothelial cells respectively (Fig. 6b-g). Similar observations were also noted in helper T cells (CD4) and cytotoxic T cells (CD8), where the frequency of CD3+CD4+ and CD3+CD8+ cell population was significantly lower in porcine kidney ST8Sia6 culture conditions (Fig. 6h-i). However, no significant difference in the frequency of helper T cells (CD4) and cytotoxic T cells (CD8) populations were observed in the porcine aorta experimental condition (Fig. 6j-k). Furthermore, the expression of activation markers such as CD69, CD25 and CD28 on T cells and NKG2D, NKp46, and DNAM1 on NK cells was significantly reduced in PBMC after interaction with ST8Sia6-overexpressing porcine cells during coculture condition. Conversely, the expression of inhibitory markers like PD-1, LAG-3 and CD106 on T cells and 2B4, CD94, CD122 and KLRG1 on NK cells was increased in ST8Sia6 co-culture condition (Fig. 6l-o, Supplementary Fig. 5d). Moreover, comparable trends were observed in helper T cells (CD4), cytotoxic T cells (CD8), and natural killer T (NKT) cells, showing a significant decrease in the frequency of activation markers such as CD69, CD25, CD28, NKG2D, NKp46, and DNAM1 expression respectively during ST8Sia6-overexpressing porcine cells during coculture conditions (Fig. 6l-o, Supplementary Fig. 5d). These results support the reduced level of cytotoxic killing seen endothelial cells expressing ST8Sia6 further demonstrating that ST8Sia6 overexpression has a dampening effect on the activation of various T cell and natural killer (NK) subsets. Additionally, these findings suggest that overexpression of ST8Sia6 in porcine cells can modulate the activation state and cytotoxic potential of human T cells and NK cells by regulating cytokine and chemokine secretion such as IFN-γ, perforin, granzyme-B, which play a vital role in immune mediated destruction for xenograft rejection. Therefore, immune cells response was further assessed using a degranulation assay, which detects the production of effector molecules including cytokines and cytolytic components following interference with ST8Sia6 overexpression in porcine kidney and aorta cells. This assessment focused on measuring CD107a expression in NK cells, T cells, and specific T cell subpopulations such as CD4+, CD8+, and CD4+CD8+ cells. The results demonstrated that ST8Sia6 overexpression in porcine cells significantly diminished CD107a expression on these cell types (Fig. 6p-s), indicating a reduction in their cytotoxic potential and secretion of cytotoxic molecules. This suggests that ST8Sia6 may play a pivotal role in modulating the immune response against xenograft transplants by attenuating the activation and cytotoxicity of human T cells and NK cells. The observed effects on T cell and NK cell activation and inhibition markers suggest that ST8Sia6 overexpression in porcine aorta cells dampen the cytotoxic potential of human immune cells, potentially contributing to improved xenograft survival. The efficacy of effector molecule suppression in porcine kidney endothelial cells was not as effective as measured in porcine aorta. However, the reduced levels of IFN-g and other regulatory markers indicate that, despite their increased presence in the kidney, these molecules seem to be ineffective in triggering a strong cytopathic effect, since there was marked differences in CTL killing of kidney endothelial.

**Figure 6.**
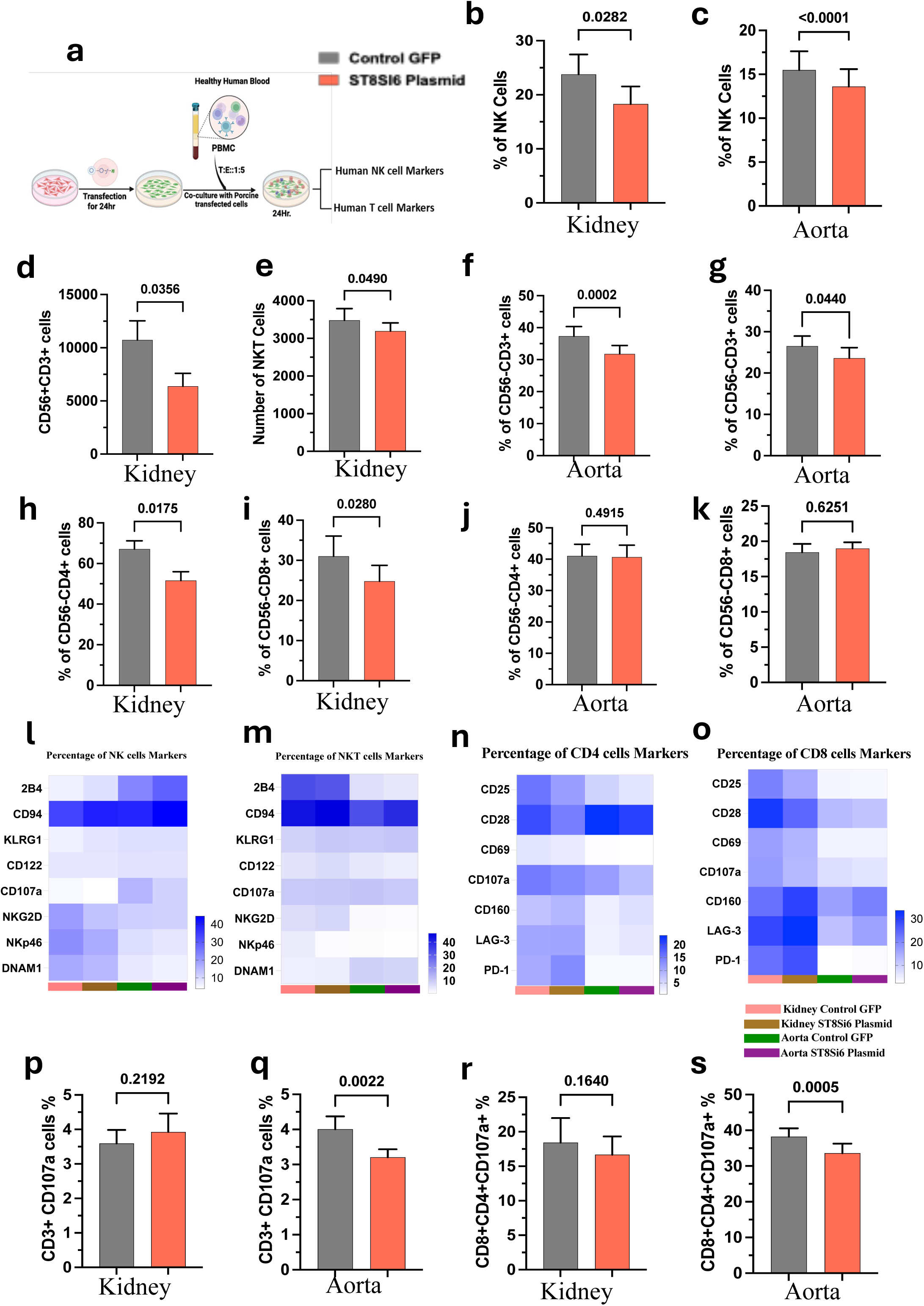
Sialic acid over expression modulates the immune cells activation and surface marker expression. a, Schematic representation of immune cell activations in-vitro experimental designed of ST8Sia6 overexpressing porcine kidney and aorta endothelial cells co-culture in presence of healthy human PBMC respectively. All the graphs represent percentage of different immune cells populations such as **b-c,** NK cells, **d-e,** NKT cells (CD56+CD3+), **f-g,** CD4 cells, **h-i,** CD-3 T cells (CD56-CD3+), **j-k,** CD8 T cells (CD56-CD8+). **l-o,** Hitmap represent expression of different activation and inactivation surface marker for NK and T cells. **p-s,** Percentage of CD3+CD107a+ cells, Percentage of CD8+CD4+ CD107a+ Cells. All graphs summarize the results from 4 donors (mean ± S.E) with 4 individual replicates. Significant difference (P<0.05) assessed by paired t-test, and the P values are represented in the graphs.

Further investigation is needed to elucidate the precise mechanisms by which ST8Sia6 modulates the xeno-immune response, and the differences observed in porcine aorta and kidney endothelial cells. These differences warrant further study to better understand how to pre-treat and maintain different xeno-transplanted organs. To assess the long-term implications of forced ST8Sia6 expression in these two tissue sources, a phosphorylation screen of immunoreceptor tyrosine-based inhibition motifs (ITIMs) and immunoreceptor tyrosine-based activation motifs (ITAMs), as well as transcriptomic analysis, was conducted.

### ST8Sia6 Overexpression in Porcine Endothelial Cells enhances ITIM expression on human immune cells from bulk peripheral blood, leading to inhibition of immune effector mechanisms

Immunoreceptor Tyrosine-based Inhibition Motifs (ITIMs) and Immunoreceptor Tyrosine-based Activation Motifs (ITAMs) are essential phosphorylation motifs that significantly influence the immune response. The dynamic interaction between ITIMs and ITAMs is crucial for maintaining immune homeostasis ^24,53^. These motifs are found in various immune receptors, including Fc receptors for antibodies, as well as certain inhibitory receptors on natural killer (NK) cells and T cells. When sialylated ligands interact with inhibitory receptors (e.g., Siglecs), ITIM-mediated signals dominate, suppressing immune cell activation. Conversely, if sialic acid structures are absent or reduced, ITAM-mediated signals are more likely to proceed unchecked, promoting immune activation ^54^. ITIMs function as inhibitory signals that effectively counterbalance the activating signals mediated by ITAMs, thereby facilitating the exploration and development of novel immunotherapies. The sialic acid linkage can modulate the binding and signaling capacity of immune receptors that contain ITIMs and ITAMs, which in turn regulates immune cell activation and exhaustion ^55^. Understanding these mechanisms could provide insights into the regulation of immune responses and potential therapeutic interventions in xenotransplantation targeting sialic acid-mediated pathways. We assessed the expression of various ITIM and ITAM motifs in immune cells from human peripheral blood after co-culturing ST8Sia6-transfected porcine aorta and kidney endothelial cells with human peripheral blood mononuclear cells (PBMCs) for 24 hours. Our aim was to determine if the phosphorylation status of ITIM and ITAM receptors reflected the immune regulatory levels observed in immune effector mechanisms from human peripheral blood immune cells incubated with ST8Sia6-expressing porcine kidney and aorta endothelial cells. Our flowcytometry estimation indicate that the expression of ITIM-containing receptors, such as TIAIT showed a strong trend in expression in immune cell populations including NK cells, NKT cells, T cells, and B cells under the porcine kidney and aorta co-culture conditions with ST8Sia6 compared to the control condition with GFP expression (Fig. 7a, Supplementary Fig. 7a, Supplementary Fig. 8a and c). In contrast, ITAM-containing receptors such as TREM showed a significant reduction in expression across various immune cell subpopulations when co-cultured with porcine kidney and aortic endothelial cells expressing the sialyltransferase ST8Sia6 (Fig. 7b, Supplementary Fig. 7b, Supplementary Fig. 8b and d). While the absolute number of inhibitory TIAIT-expressing immune cells is significantly higher in the kidney and aorta under ST8Sia6 transfected conditions (see Supplementary Fig. 8a-b). Conversely, the absolute number of TREM-expressing immune cells is significantly lower in the same conditions (see Supplementary Fig. 8c-d). This suggests that sialic acid linkages may play a crucial role in balancing activating and inhibitory immune signaling by modulating targeted sialic acid-Siglec interactions on immune cells during cell-cell interaction. To further quantitate the expression levels of ITIM and ITAM-associated proteins under ST8Sia6 overexpression conditions, we performed western blot analysis using PBMC lysates derived from a porcine aorta and kidney endothelial co-culture experiment. Anti-DAP12 and anti- TREM2 primary antibodies were used to detect ITAM-associated proteins, while anti-TIGIT, anti-SHP-2, anti-pSHP-2, and anti-SHP-1 primary antibodies were employed for ITIM-associated protein detection. The results revealed a reduced expression of DAP12 and TREM2 in human PBLs containing immune cell lysates obtained from ST8Sia6-transfected porcine kidney cells co-conditions compared to the group transfected with control-GFP (Fig. 7c) and similar observation also detected into ST8Sia6-transfected porcine aorta endothelial cells co-culture conditions (Fig. Supplementary 7c). Furthermore, western blot analysis showed that the expression levels of ITIM-associated proteins, such as TIGIT, SHP-2, p-SHP-2, and SHP-1, under similar conditions were highly express in PBL immune cells co-cultured with ST8Sia6-transfected porcine aorta as well as kidney endothelial cells (Fig. 7d, and Supplementary 7d). Taken together the above findings suggest that ST8Sia6 over expression in porcine endothelial cells modulates the expression of ITIM-associated proteins in co-cultured immune cells. The increased levels of ITIM proteins, such as TIGIT, SHP-2, and SHP-1, indicate a potential regulatory role of ST8Sia6 in dampening immune cell activation and promoting an anti-inflammatory state, corroborating the results shown above, thus supporting the immune suppressive role ST8Sia6 has on effector mechanism expression. Further investigation is needed to elucidate the underlying mechanisms by which ST8Sia6 influences the ITIM/ITAM balance in the endothelial-immune cell crosstalk.

**Figure 7.**
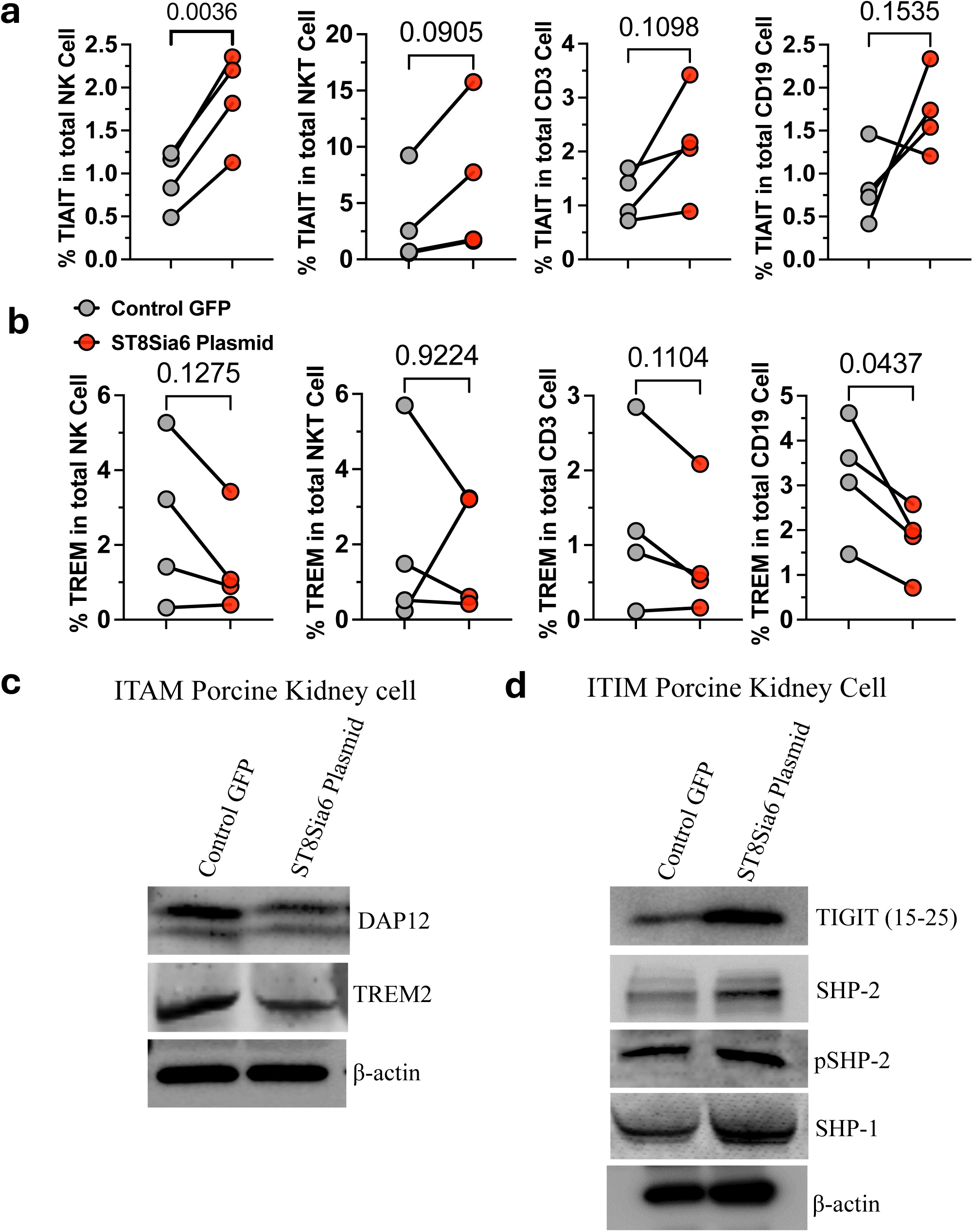
The overexpression of sialic acid in porcine kidney endothelial cells primarily drives immune regulation through ITAM/ITIM signaling pathways in various immune cells. **a**, percentage of TIAIT a tyrosine-based inactivation motif expression in different immune cells with in ST8Sia6 and Control GFP expressing group. **b,** Proportion of TREM a tyrosine-based activation motif expression across different immune cell types in the ST8Sia6 group compared to the Control GFP-expressing group. **c-d,** Assessment of ITAM and ITIM protein expression in PBMC lysates from the porcine kidney endothelial co-culture condition probe with anti-DAP12, anti-TREM2 primary antibody for detection of ITAM and anti TIGIT, anti-SHP-2, anti pSHP-2, anti- SHP-1, primary antibody for ITIM detection. Graphs show data from 4 donors (mean ± S.E) in average 4 replicates. Significant differences (P<0.05) were determined by paired t-test, and P values are displayed in the graphs.

### Transcriptomic alteration induced by sialic acid overexpression in porcine kidney endothelial cells

To investigate the molecular changes associated with ST8Sia6 overexpression in porcine kidney endothelial cells, we performed RNA sequencing on four independent experimental setups comparing control GFP-expressing cells to ST8Sia6-overexpressing cells. Differential gene expression analysis using a stringent criteria (false discovery rate (FDR) <0.05 and a log2fold change ≥|1.5) identified a total of 209 differentially expressed genes (DEGs). Among these, 64 genes were upregulated and 145 were downregulated in ST8Sia6-overexpressing cells compared to the control (Fig. 8a). To gain insight into the biological implications of these DEGs, we conducted gene set enrichment analysis (GSEA). This analysis revealed that ST8Sia6 overexpression affected several key biological processes, including cellular processes, immune regulation, immune cell-mediated response, metabolic regulation, multicellular biological processes, cellular interactions, and immune cell functional activity ^56^. Notably, in response to ST8Sia6a overexpression, B cell receptor signaling pathway, T cell receptor signaling pathway, and NK cell-mediated cytotoxicity, leukocyte transendothelial migration, ABC transporters, phosphatidylinositol signaling, primary immunodeficiency, and complement activation were significantly downregulated (Fig. 8b, supplementary Fig. 10).

**Figure 8.**
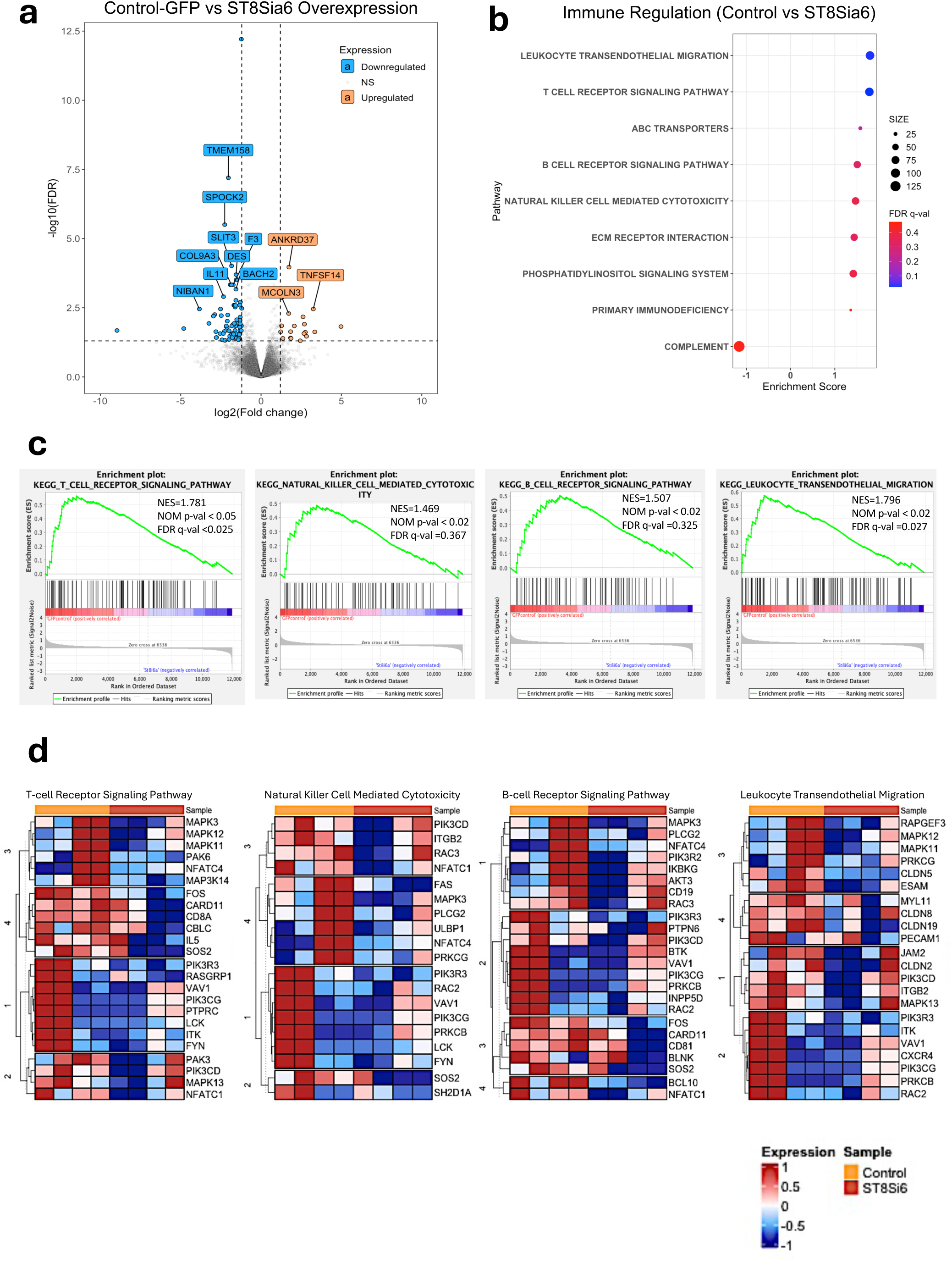
Overexpression of ST8Sia6 in porcine kidney endothelial cells effectively modulates transcriptomic changes in immune regulatory pathways. **a.** Volcano plots of differentially expressed genes (DEGs) between control porcine kidney cells and sialic acid overexpressing porcine kidney cells. The orange and blue dots represent upregulated and downregulated DEGs respectively with an FDR ≤ 0.05 and a log2 (FoldChange) of ≥|1.5|. The dashed horizontal line indicates the FDR threshold of 0.05, while the vertical dashed lines represent the log2 (FoldChange) thresholds of ≥|1.5 |. **b.** Dot plot for enriched pathways involved in immune regulation, comparing control-GFP and ST8Sia6 overexpressing porcine kidney endothelial cells. Dot size is proportional to normalized enrichment score (NES) and P-values are color-coded according to the color scale. **c.** Enrichment plots for top five **e**nriched pathways involved in T-cell receptor, Natural Killer cell, B-cell receptor and leukocyte transendothelial migration showing the profile of the running ES Score and positions of gene set members on the rank-ordered list. **d**. Heat map of the top 30 marker genes for each phenotype involved in immune regulation comparing ControlGFP (Yellow, left three columns) vs. ST8Sia6 overexpress (Orange, right three column). Red, upregulated; Blue, downregulated genes.

Heatmap analysis of significant genes revealed distinct expression patterns downstream of immune-mediated modulatory pathways. Where ST8Sia6-overexpressing cells led to downregulation in immune activated gene signatures. These included genes associated with immune activation and complement activation, such as Mitogen-activated protein kinase (MAPK), Interleukin 5 (IL5), CD81, Ras-related C3 botulinum toxin substrate 3 (RAC3), P21-activated kinase 3 (PAK3), BCL10 and Fas cell surface death receptor (FAS) (Fig. 8d). Additionally, genes linked to ABC transporter signaling pathways, hallmark compliment pathway related genes exhibited significant downregulation under ST8Sia6 overexpression conditions (Supplementary Fig. 10b). Together, these findings provide a comprehensive understanding of immune-mediated modulatory pathways affected by ST8Sia6 overexpression, potentially reducing immune hyper- reactivity during xenotransplantation. This regulation of immune signaling pathways by sialic acid may have significant implications for improving transplant tolerance and reducing rejection responses in xenogeneic transplantation models.

We further investigated the DEGs associated with the metabolic signature to identify key pathways affected by ST8Sia6 overexpression. The dot plot analysis revealed that the top five metabolic gene sets were significantly upregulated in ST8Sia6-overexpressing cells compared to control-GFP cells and included oxidative phosphorylation, glycolysis, fatty acid metabolism, xenobiotic metabolism, hedgehog signaling (Fig. 9a). In contrast, gene sets associated with basal transcription factor, arginine and proline metabolism, pyruvate metabolism, and pyrimidine metabolism were significantly enriched in control-GFP porcine kidney endothelial cells compared to ST8Sia6-overexpressed porcine kidney endothelial cells (Fig 9b-c, Supplementary Fig. 11a). A hierarchical heatmap displaying the top 30 marker genes from various enriched pathways highlighted distinct expression patterns in ST8Sia6-overexpressing cells (Fig 9b, Supplementary Fig. 11b). Clustering analysis of DEGs mapped to KEGG pathways revealed distinct functional modules, displaying dense interactions among proteins involved in energy metabolism and cellular growth regulation (Supplementary Fig. 11c). These findings highlight the significant impact of ST8Sia6 overexpression in porcine kidney endothelial cells, particularly its ability to modulate the expression profiles of specific ligand-associated genes. Notably, the affected gene signatures are enriched in pathways related to immune signaling, complement activation, cellular metabolism, and differentiation. Together, these results point to a pivotal role for ST8Sia6 in regulating immune functions and developmental processes.

**Figure 9.**
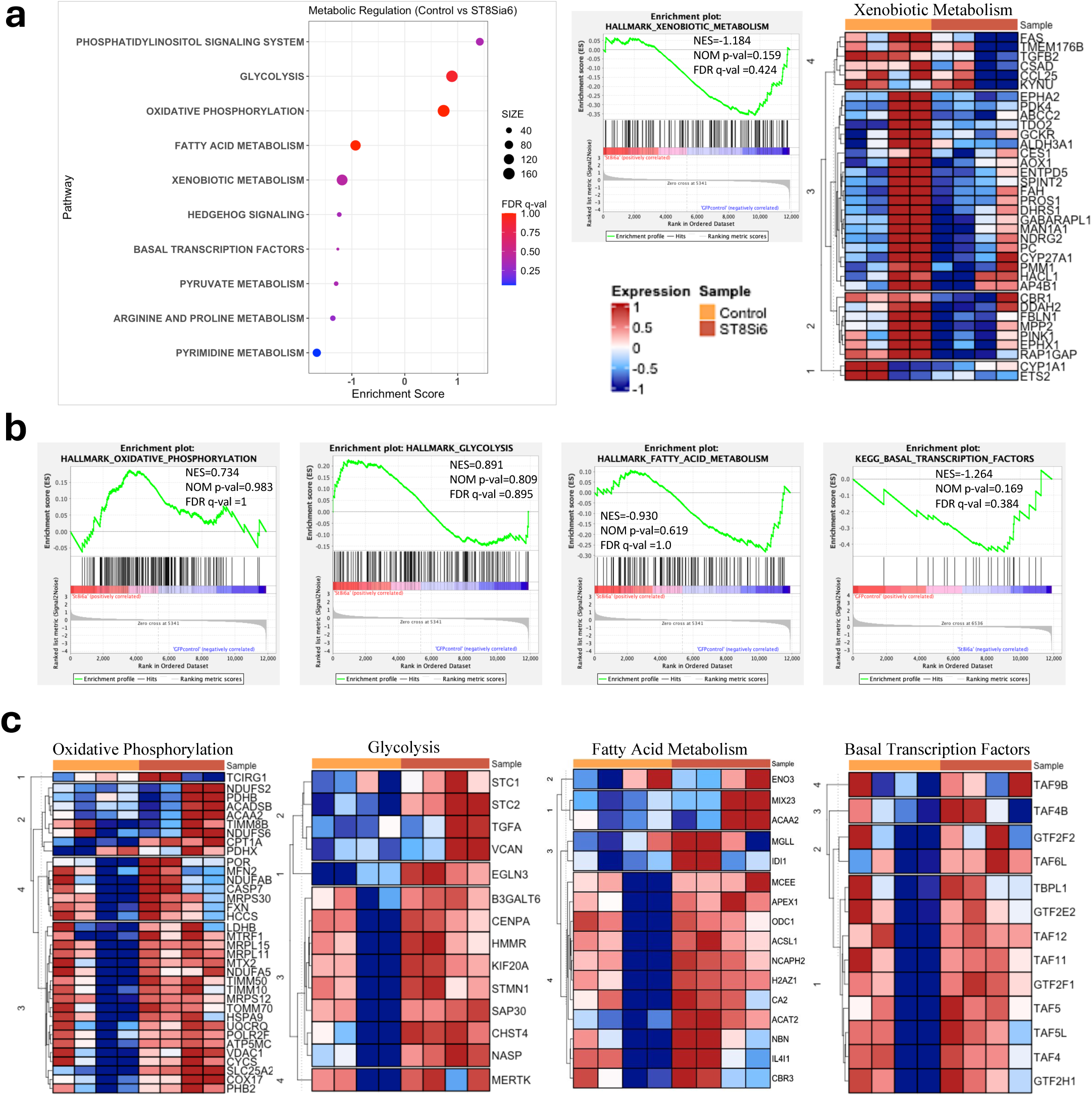
Overexpression of ST8Sia6 in porcine kidney endothelial cells effectively modulates transcriptomic changes in metabolic pathways. **a.** Dot plot for enriched pathways involved in metabolic regulation, comparing control-GFP and ST8Sia6 overexpressing porcine kidney endothelial cells. Dot size is proportional to NES and P-values are color-coded according to the color scale. **b.** Enrichment plots for top five **e**nriched pathways involved in adaptive immune system pathways, such as oxidative phosphorylation, glycolysis, fatty acid metabolism, and basal transcription factors. **c.** Heat map of the top genes involved for metabolic pathways for each phenotype in the comparison of ControlGFP (Yellow, left three column) vs. ST8Sia6 overexpress (Orange, right three column). Red, upregulated; Blue, downregulated genes.

### Transcriptomic changes induced by sialic acid overexpression in porcine kidney endothelial cells co-cultured with human PBMCs

To understand the biological functions of DEGs associated with ST8Sia6 overexpression in porcine kidney endothelial cells upon co-culture with human PBMCs, transcriptomic analysis was performed on four independent experimental conditions, including control GFP-expressing cells and ST8Sia6-overexpressing cells. A total of 3,839 DEGs were identified using an FDR threshold of <0.05 and a log2fold change ≥|1.5|. A comparative analysis between GFP-Control vs GFP- Control porcine cells from coculture with human PBMC revealed 3849 genes were upregulated (Fig. 10a), interestingly in ST8Sia6-overexpressing cells vs ST8Sia6-overexpressing porcine cells from coculture revealed 1,599 upregulated and 2,240 downregulated genes as visualized in the volcano plot (Fig. 10b). Additionally, comparison of gene expression between the GFP-Control and ST8Sia6-overexpressing groups under cell–cell interaction conditions revealed that only 143 genes were upregulated, while 580 genes were downregulated (Fig. 10c). Furthermore, GSEA of normalized gene expression using KEGG and hallmark gene expression analysis identified several pathways significantly enriched in control GFP-expressing cells compared to ST8Sia6- overexpressing cells during co-culture. These included receptor interaction pathways, NOD like receptor signaling TGF beta signaling pathway (Fig. Supplementary Fig. 10d-e). These findings highlight the differential regulation of key signaling pathways in response to ST8Sia6 overexpression during interactions with human PBMCs.

**Figure 10.**
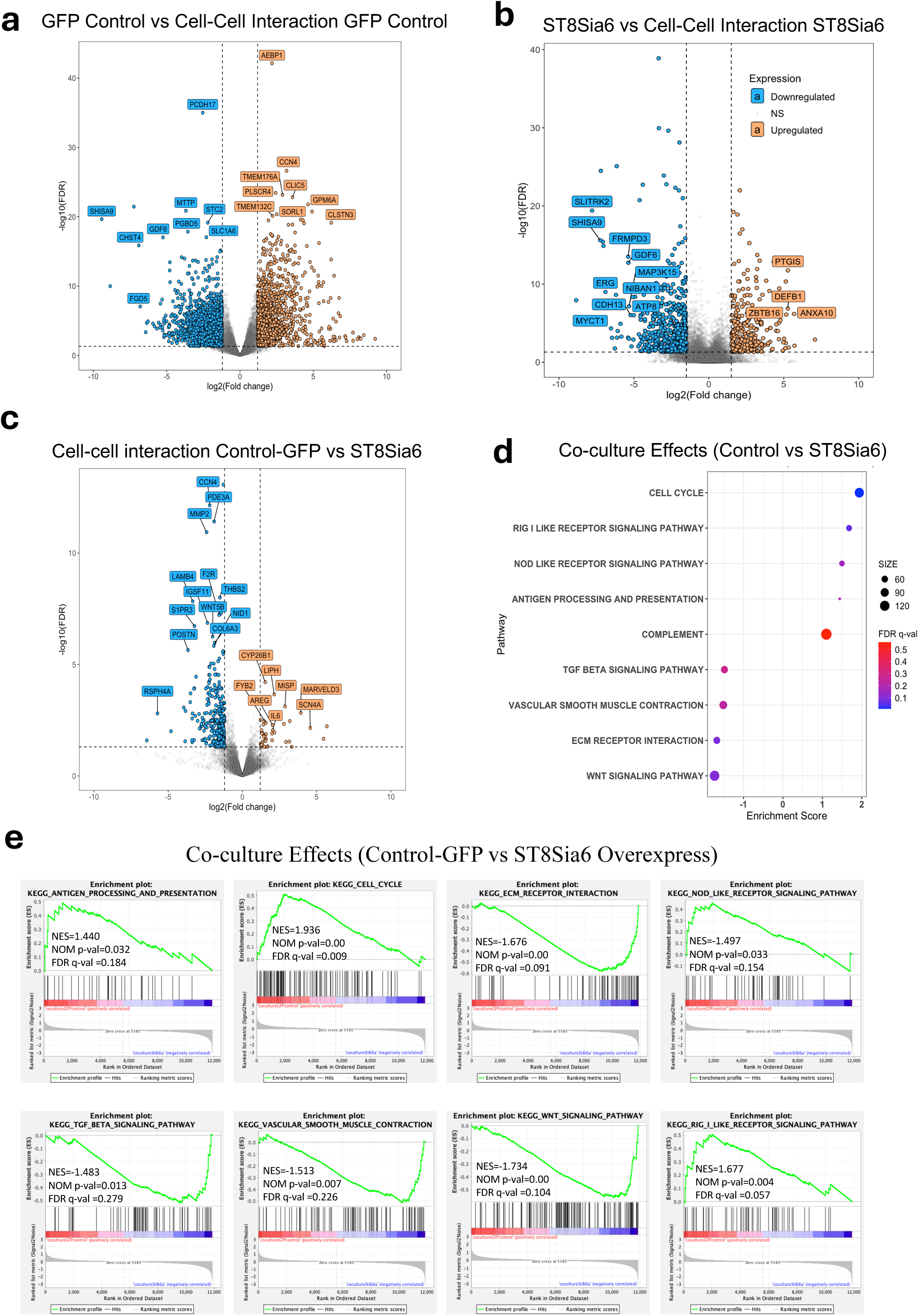
Volcano plot of DEGs comparing **a.** GFP control vs GFP control co-cultured with human PBLs. **b.** ST8Sia6 overexpressed porcine kidney compared to ST8Sia6 overexpressed porcine kidney cell co-cultured with human PBLs. **c.** GFP control co-cultured with human PBLs vs ST8Sia6 overexpressing porcine kidney cell co-cultured with human PBLs. Orange, upregulated and Blue, downregulated genes. **d.** Dot plot for enriched pathways involved in different cellular and immune pathway in comparison with control-GFP and ST8Sia6 over expressing porcine kidney endothelial cells. Dot size is proportional to NES and P-values are color-coded according to the color scale. **e.** Enrichment plots for top eight **e**nriched pathways involved in different cellular and immune pathways showing the profile of the running ES Score and positions of gene set members on the rank-ordered list.

## Discussion

The global shortage of organs from deceased human donors for patients with end-stage organ failure remains critical. Xenotransplantation, particularly using gene-edited pig organs, is considered the most promising alternative ^57^. Pigs have been identified as the primary source for clinical xenotransplantation, with recent research focusing on pig-to-human transplantation despite significant immunological barriers ^58^. Advancements in genome editing, especially CRISPR-Cas9 technology, have led to significant progress in xenotransplantation ^8,59^. These advancements include the elimination of carbohydrate xenoantigens from pig cells and the FDA approval of GalSafe pigs ^8^. These changes have set a new standard in preventing the recognition and binding of preformed human antibodies, thereby reducing the risk of hyperacute rejection. This is achieved by knocking out the (1,3) galactosyltransferase (a1,3GT) gene, which removes glycoconjugates from the porcine cell surface, delaying xenograft rejection ^60^.

Our understanding of sialic acid linkage differences in various species has improved, revealing how these differences modulate immune tolerance and play a crucial role in self/non-self-recognition. Additionally, we have gained insights into how disease states like cancer use these sialic acid linkages to evade immune recognition by tumor-infiltrating immune cells, as well as the essential role these sialic modifications play in ensuring successful fetal development during pregnancy. Tumor cell sialoglycans promote survival and growth by evading immunosurveillance ^42^. While tumors are known to incorporate more sialic acids in surface glycans, the role of individual sialyltransferases and their mechanisms in cancer remain largely unexplored. Exploiting these methods of natural immune inhibition prompted us to use this strategy to shroud porcine tissue in a human sialic acid linkage on the porcine cell surface by force expressing the human enzyme ST8Sia6-sialyltransferase in porcine kidney and aorta cells.

Our study first emphasized the significant role ST8Sia6 in modulating sialylation patterns and its impact on immune interactions in the widely used B16F murine melanoma cell line. The upregulation of ST8Sia6 in various human cancers, as supported by publicly available databases, suggests its involvement in tumor progression and immune evasion. Our engineered B16-F10 and B16-Ova melanoma cell lines with stable ST8Sia6 expression provided a well-described model system to test the efficiency of transient overexpression strategies in glycosylation-related modifications and their effects on immune function.

The findings from our study provide valuable insights into the role of 2,8-linked sialic acid in modulating immune interactions, particularly in the context of xenotransplantation. The overexpression of the ST8Sia6 gene in porcine kidney and aorta endothelial cells resulted in significant expression of 2,8-linked sialic acid, that modification plays a key role in immune down regulations ^30^. By transfecting these porcine endothelial cells with ST8Sia6, evidence demonstrated a clear, receptor-specific interaction with Siglec-E and Siglec-9 receptors showed significantly higher binding to ST8Sia6-transfected porcine kidney and aorta cells compared to control-GFP conditions, which is strongly support our hypothesis (Supplementary Fig. 4a-d) ^61^. These Siglecs are known to be involved in immune signaling, and their binding to the 2,8-linked sialic acid confirms its potential to modulate immune responses.

Our data demonstrates that ST8Sia6 expression significantly increased the binding frequency of Siglec-E to B16F10-melanoma cells, as well as the porcine kidney and aorta cells indicating the formation of α2,8-linked disialic acid structures, which are known ligands for Siglec-E supports the hypothesis that ST8Sia6-mediated modifications play a role in immune modulation, potentially contributing to immune evasion mechanisms demonstrated in our results The lectin binding assays in B16-melanoma cells using PNA further elucidated the impact of ST8Sia6 expression on O-linked glycan accessibility. The observed reduction in PNA binding in ST8Sia6-expressing B16-F10 cells indicates that α2,8-linked sialylation interferes with O-glycan recognition, potentially altering cell surface interactions with immune cells ^62^. The increased binding of Siglec-E and Siglec-9 to ST8Sia6-transfected porcine kidney and aorta cells is of particular significance, since engagement of these receptors, by their ligands, mediates inhibitory signals that dampen immune responses ^63,64^. In the context of xenotransplantation, we provide strong evidence that this interaction reduces the likelihood of optimal immune recognition leading to rejection, as it limits the activation of human innate and adaptive arms of the immune system including, T cells, NK cells, macrophages and activation of other immune components, like complement cascade that typically become activated to foreign porcine tissues ^65,66^. In contrast, control cells displayed higher PNA binding, suggesting an abundance of unmodified O-linked glycans available for recognition. Another important finding from this study is the relatively high transfection efficiency (55% for porcine kidney cells and 50% for aorta cells), which ensures that enough cells express the transgene. The detection of ST8Sia6 protein expression in the transfected cells further supports the successful generation of porcine cells with the sialic acid modification. These findings collectively suggest that ST8Sia6 overexpression leads to a shift in glycan expression, influencing immune cell interactions, hyperacute recognition, and immune evasion mechanisms ^30,67^. Therefore, our recent findings suggest that incorporating 2,8-linked sialic acid on porcine endothelial cells could potentially improve the success rates of xenotransplantation by lowering immune activation and promoting immune tolerance. Interestingly, no significant binding was observed with Siglec-10, indicating the specificity of the interaction between 2,8-linked sialic acid and Siglec-E and Siglec-9. This specificity is important, as it highlights the potential for designing targeted strategies to manipulate immune interactions in xenotransplantation. The absence of Siglec-10 binding also underscores the complexity of Siglec-receptor interactions and suggests that not all Siglec receptors are equally involved in the immune response to 2,8-linked sialic acids ^24,68,69^. Further studies we explore the exact mechanisms behind these interactions, the broader impact of 2,8-linked sialic acid on immune responses.

Our results demonstrate the importance 2,8-sialic acid linkage expression, from ST8Sia6 forced expression in negatively regulating immune function, particularly its impact on immune cell-mediated cytotoxic and innate immune components in xenotransplantation rejection. Our findings reveled that, ST8Sia6 expression in B16-F10 and B16-Ova cells suppressed CD8+ T cell-mediated cytotoxicity, as evidenced by reduced killing efficiency and decreased IFN-γ production ^70,71^. Furthermore, the downregulation of key activation markers, CD69 and LAMP-1, suggests that 2,8-linked sialic acid may suppress T cell activation through immune checkpoint-like mechanisms, contributing to protect hyperacute rejection of xenograft in clinical aspects ^72^. In the context of xenotransplantation, ST8Sia6-expressing porcine kidney and aorta endothelial cells exhibited significantly reduced susceptibility to cytotoxic killing by human PBMCs, accompanied by decreased secretion of IFN-γ, perforin, and granzyme-B. our results support that ST8Sia6 expression modulates immune responses by altering cytokine secretion, immune cell activation, mitigating immune-mediated xenograft rejection ^30^. Collectively, our study highlights, the ST8Sia6- Siglec-9 axis as a crucial immune regulatory pathway, suggesting its potential as a therapeutic target for improving xenograft acceptance by suppressing immune-mediated rejection.

Furthermore, our result demonstrates that, ectopic expression of ST8Sia6 in porcine endothelial cells has been shown to mitigate the Instant Blood-Mediated Inflammatory Reaction (IBMIR), a significant challenge in xenotransplantation. IBMIR involves immune activation, complement cascade, coagulation, and inflammatory responses, which often lead to rapid organ rejection ^46,73^. Our study demonstrates that, a2,8 sialic acid linked porcine kidney and aorta endothelial cell interaction with human immune cells significantly reduced complement activation and coagulation (e.g. decreased C3a, C4b, C5b9 production, and the TAT complex formation). This suggest that a2,8-linked sialic acid overexpression could enhance xenograft survival and function by prevent the complement mediated inflammatory immune response^74^. Furthermore, a2,8-linked sialic acid in conjunction with complement regulators like Decay-Accelerating Factor (DAF) and CD46, appears to inhibit the formation of the membrane attack complex and stop further complement activation ^75^. Collectively our findings underscore the ability of ST8Sia6 to overcome IBMIR and improve xenotransplantation outcomes, although additional research is needed to fully assess its effects in preclinical models.

Our study provides compelling evidence that, ST8Sia6 overexpression in porcine endothelial cells modulates immune responses by regulating the activation and inhibition of both innate immune as well as cellular specific mechanisms, such as inhibition of complement activation and regulation of human T cells and NK cells. Friedman et al. (2022) demonstrated that ST8Sia6 knockout mice exhibited enhanced anti-tumor immunity, characterized by increased activation of macrophages and dendritic cells, as well as a shift in regulatory T cells towards a more inflammatory phenotype ^76^. Our data corroborate these findings as we also see a significant reduction in the frequency of CD3+ T cells, CD56+ NK cells, and CD3+CD56+ NKT cells following co-culture with ST8Sia6-expressing porcine kidney and aorta endothelial cells suggests that ST8Sia6 expression influences both adaptive and innate immune responses. These findings align with previous reports emphasizing the critical role of NK cells in the initial response against xenograft rejection, as well as the importance of T cell-mediated cytotoxicity in shaping immune responses to transplanted tissues ^77^. Significant downregulation of key activation markers (CD69, CD25, CD28 on T cells; NKG2D, NKp46, DNAM1 on NK cells) was observed, reinforcing the notion that ST8Sia6 overexpression may suppress immune cell activation. Conversely, the upregulation of inhibitory markers (PD-1, LAG-3, CD106 on T cells; 2B4, CD94, CD122, KLRG1 on NK cells) highlights the potential immunosuppressive role of ST8Sia6 in xenotransplantation. ^51,78^. Our recent studies have investigated the role of ST8Sia6 overexpression in modulating immune responses, particularly its impact on CD107a expression in various immune cell subsets. CD107a, also known as lysosomal-associated membrane protein-1 (LAMP-1), is commonly used as a marker for degranulation in cytotoxic T cells and natural killer (NK) cells. In a study by Friedman et al., findings suggest that ST8Sia6-mediated modulation of Siglec-E could potentially influence degranulation processes in other immune cells, such as T cells and NK cells. ^30^. The ST8Sia6 overexpression significantly diminished CD107a expression across multiple immune subsets, including CD4+, CD8+, and CD4+CD8+ T cells, as well as NK cells. Since CD107a is a critical marker of cytotoxic granule release, its reduced expression indicates a functional impairment in the cytotoxic potential of these immune cells ^79^. The concurrent decrease in Hu- IFN-γ, Hu-perforin, and Hu-granzyme-B secretion through immune cells during co-culture, further suggests that ST8Sia6 overexpression attenuates immune effector functions necessary for xenograft rejection. These results suggest that reducing immune effector mechanisms could help prevent chronic rejection in xenotransplants. Chronic rejection is driven by the gradual loss of tolerance and the activation of T cells specific to xenoantigens, leading to graft failure. By impeding these responses early, as our results indicate, we may lower the frequency of xeno - targeted antigens, prolong tolerance, and, when combined with standard immunosuppression, better manage chronic rejection pathways. Additionally, study by Chou-Yuan et al., bioinformatics analyses have identified ST8Sia6 as a potential therapeutic target in colon cancer, with its expression affecting immune cell infiltration and responses to immunotherapy. These findings further support the notion that ST8Sia6 plays a role in modulating immune cell functions, possibly including degranulation processes marked by CD107a expression ^56^. Taken together, these results suggest that ST8Sia6 expression modulates immune responses by altering cytokine secretion, reducing activation markers, and increasing inhibitory signals, fostering a tolerogenic environment for xenograft survival. Targeting the ST8Sia6-Siglec-9 axis could represent a novel approach to mitigate immune rejection in xenotransplantation.

Moreover, exploring the influence of porcine cells expressing sialic acid linkage on ITAM (Immunoreceptor Tyrosine-based Activation Motif) and ITIM (Immunoreceptor Tyrosine-based Inhibition Motif) expression in immune cells is of particular interest in xenograft rejection. ITAM and ITIM are critical in modulating immune cell activation and inhibition, respectively ^53^. Our findings suggest that the expression of sialic acid linkages on porcine cells may influence ITAM-associated activation pathways, leading to decreased immune responsiveness. Concurrently, our study demonstrated that overexpression of ST8Sia6 in tumor cells increased α2,8-linked disialic acids, which engaged Siglec-E on murine immune cells. This interaction activated ITIM domains, leading to immune suppression and which reinforces our hypothesis that these linkages play a critical role in ITIM mediated immune inhibition during xenotransplantation ^30,80^. Further, in our studies we investigate the specific ST8Sia6-overexpressing porcine endothelial cells significantly influence the balance between ITIM and ITAM signaling in immune cells. The increased expression of ITIM-associated receptors such as TIAIT and inhibitory proteins including TIGIT, SHP-2, and SHP-1 suggests a shift toward an immunosuppressive phenotype. Friedman et al. demonstrated that ST8Sia6 expression on tumor cells is linked to altered macrophage polarization toward the immunosuppressive M2 phenotype. This polarization involves the upregulation of immune modulators like arginase, a process dependent on Siglec interactions. These findings strongly support our results, indicating that the downregulation of ITAM-associated receptors, such as TREM, along with their signaling intermediates, including DAP12 and TREM2, underscores a reduction in immune activation ^30,81^. This dynamic interplay indicates that sialic acid linkages may modulate the immune response by favoring inhibitory over activating signals, potentially contributing to immune tolerance in xenotransplantation. Western blot analysis confirmed these findings, showing a significant increase in ITIM-associated proteins, including phosphorylated SHP-2 and SHP-1, while ITAM-related proteins such as DAP12 and TREM2 were reduced under ST8Sia6-overexpressing conditions. This suggests that ST8Sia6 might facilitate immune regulation by shifting the balance toward inhibitory signaling, ultimately reducing immune-mediated xenograft rejection. Taken together, these findings indicate that ST8Sia6 overexpression in porcine endothelial cells may modulate the expression of ITIM/ITAM-associated proteins in co-cultured immune cells. The increased levels of ITIM proteins, including TIGIT, SHP-2, and SHP-1, suggest that the interaction between a2,8-Sialic acid and Siglecs on the porcine tissue is functioning to regulate the self-non-self-recognition of the a2,8 sialic acid expressing-porcine tissue, treating it like normal human tissue. This is what we expected, and we were surprised that we had this type of impact with only the expression of this one gene in these otherwise wild type porcine endothelial cells. It is tempting to speculate that in conjunction with the triple knock-out” (TKO) pigs we would see an even better level of tolerance to these tissues leading to a regulatory role in suppressing immune cell activation, fostering an anti-inflammatory state, and reduce chronic rejection. Studies by Noguchi et al., showed that porcine aortic endothelial cells transfected with human TIGIT (PAEC/TIGIT) have revealed significant immunosuppressive effects. Additionally, a notable increase in phosphorylated SHP-1 was observed in M1 macrophages co-cultured with PAEC/TIGIT showed reduced cytotoxicity and decreased expression of pro-inflammatory cytokines, such as TNFα, IL-1β, and IL-12 ^82^. By expressing ST8sia6 in these porcine endothelial cells we provide an all-encompassing level of inhibitory signals maintaining tolerance to porcine tissue expressing ST8sia6. Collectively, our findings provide compelling evidence that ST8Sia6 expression has the potential to influences the ITIM/ITAM balance in the endothelial-immune cell crosstalk. Moreover, investigating the clinical and therapeutic potential of targeting sialic acid-mediated pathways could open new approaches to enhancing xenograft survival and mitigating immune rejection.

Furthermore, our study reveals that ST8Sia6 overexpression in porcine kidney endothelial cells drives significant transcriptomic changes, particularly in immune modulation and metabolic reprogramming. Differential analysis identified 64 upregulated and 145 downregulated genes, highlighting ST8Sia6’s role in immune regulation. Notably, immune-related pathways, including B cell and T cell receptor signaling and NK cell-mediated cytotoxicity, were markedly downregulated. This supports ST8Sia6’s potential to modulate immune responses by altering cytokine secretion and immune cell activation, which may help mitigate immune-mediated xenograft rejection, and reduce chronic rejection ^83^.

Gene Set Enrichment Analysis (GSEA) confirmed enrichment of immune activation pathways in control-GFP cells, while hierarchical clustering showed suppression of key immune signaling molecules such as MAPK, IL5, CD81, and FAS in ST8Sia6-overexpressing cells. This immunosuppressive effect may improve transplant tolerance by reducing immune hyper-reactivity ^42^. Beyond immune modulation, ST8Sia6 overexpression reprogrammed metabolic pathways, enhancing oxidative phosphorylation, glycolysis, and fatty acid metabolism while downregulating nucleotide biosynthesis and protein turnover. Enhancing mitochondrial function through metabolic preconditioning or mitochondrial-targeted antioxidants can improve graft survival ^84,85^. The upregulation of pyrimidine metabolism in ST8Sia6 porcine kidney endothelial cells highlights the importance of pyrimidine metabolism in cell proliferation and tissue repair after transplantation. Xenografts depend on rapid DNA and RNA synthesis for survival, making pyrimidine biosynthesis crucial for graft adaptation. While de novo synthesis is essential during high-demand periods, the salvage pathway ensures the continuous supply of nucleotides ^86^. The reduction of the xenobiotic metabolism pathway due to ST8Sia6 overexpression could potentially decrease immune rejection of transplanted tissues by altering the biochemical pathways usually targeted by the immune system, thereby enhancing graft survival ^87^. Our findings highlight ST8Sia6 as a key regulator of immune suppression and metabolic adaptation, with potential applications in reducing immune rejection in xenotransplantation. Further studies, including functional assays on immune interactions and metabolic flux analysis, will be essential for validation. Taken together, our study highlights ST8Sia6 as a critical modulator of immune and metabolic processes, with significant implications for xenotransplantation and immune regulation. The ability of ST8Sia6 to attenuate immune activation while concurrently reshaping metabolic pathways underscores its potential as a target for therapeutic interventions aimed at improving transplant tolerance and cellular resilience.

## Conclusion

Xenotransplantation presents a promising solution to the global organ shortage, with gene-edited pig organs emerging as the most viable alternative to human donor organs. However, overcoming immunological barriers remains a critical challenge. This study highlights the role of ST8Sia6- mediated α2,8-linked sialylation in modulating immune interactions, potentially enhancing xenograft survival. The overexpression of ST8Sia6 in porcine endothelial cells led to increased binding of Siglec-E and Siglec-9, which are known to mediate inhibitory immune signaling. This interaction may help suppress immune activation, reduce cytotoxicity, and promote immune tolerance, thereby mitigating xenograft rejection. Furthermore, ST8Sia6 expression influenced key immune pathways by shifting the balance between activation and inhibition, as evidenced by reduced ITAM signaling and increased ITIM-associated proteins. The observed downregulation of T cell and NK cell activation markers, along with decreased IFN-γ, perforin, and granzyme-B secretion underscores its immunosuppressive potential in xenotransplantation. Additionally, ST8Sia6 expression mitigated IBMIR, a major cause of early xenograft failure, by reducing complement activation and coagulation responses. These findings suggest that incorporating α2,8-linked sialic acid on porcine endothelial cells can improve xenograft acceptance by reducing immune activation and promoting tolerance. The ST8Sia6-Siglec-9 axis emerges as a key regulatory pathway that could be targeted to enhance xenotransplantation outcomes. Future studies should further explore the molecular mechanisms underlying sialic acid-mediated immune modulation to develop optimized strategies for clinical application in xenotransplantation.

## Materials and Methods

### Cell line culture

The B16-F10 and B16-OVA mouse melanoma cell lines were obtained from Dr. Amanda Poholek at UPMC Children’s Hospital, Pittsburgh University, PA, with a passage number of P-3 (ATCC B16-F10 Catalog CRL-6475, B16Ova Catalog CRL-21113 respectively). Both cell lines were screened for mycoplasma contamination using a mycoplasma detection kit (Catalog no. 13100- 01, Southern Biotech). Cells are cultured in DMEM (Catalog no. 11960044, Gibco) supplemented with 10% FBS (catalog no. SH30910.03, HyClone), 1% penicillin/streptomycin solution (catalog no. 15140-122, Gibco), 1% L-glutamine 200 mmol/L (catalog no. 25030-081, Gibco), 1% sodium pyruvate (catalog no. 25-000-CI, Corning), 1% HEPES (1 mol/L; catalog no. 15630-080, Gibco), 1% MEM nonessential amino acids (catalog no. 25-025-CI, Corning), and 0.1% 2- mercaptoethanol. The cells were maintained at 37°C in 5% CO2 and passaged using 0.05% Trypsin-EDTA (catalog no. 25200-056, Gibco) every 3 days. For optimal performance, cells were passaged no more than 5 times before new stocks were prepared.

### Porcine primary endothelial cell culture

The porcine primary kidney and aorta endothelial cell lines were purchased at passage P-3 and P-2 respectively from Cell Biologics (Catalog Nos. P-6014 and P-6052, respectively). The cells were cultured in tissue culture flasks pre-coated with gelatin solution (Catalog No. Cell Biologics 6950) and maintained by using Cell Biologics complete endothelial cell medium kit (Catalog No. M1168). This medium includes 0.5 mL VEGF, 0.5 mL ECGS, 0.5 mL Heparin, 0.5 mL EGF, 0.5 mL Hydrocortisone, 5.0 mL Antibiotic-Antimycotic Solution, and 25.0 mL FBS in 500 mL of basal media. The cells were kept at 37°C in 5% CO2 and passaged using 0.05% Trypsin-EDTA (Catalog No. 25200-056, Gibco). At passage 5, cells were harvested from the flasks and cryopreserved in vials, with each vial containing 0.5 × 10^6 cells per mL.

### Transfection of cell lines

Both ST8Sia6-GFP-containing (Custome made Vactor Builder, ID VB900025-4381xuz) and empty vector control-GFP (addgene #32396) plasmids were used for transfection into the cell lines mentioned above. Approximately 18-24 hours prior to transfection, around 100,000 cells were plated per well in 12-well plates, and about 50,000 cells were plated per well in 24-well plates to achieve 50-70% confluence the following day. Transfections were carried out using Promega ViaFect™ Transfection Reagent (Catalog No. E4982) in Opti-MEM™ Reduced Serum Medium (Gibco, Catalog No. 31985070), following the manufacturer’s protocol. For the primary porcine endothelial kidney and aorta cell lines, the Lipofectamine LTX reagent (Catalog No. 15338100) was employed, and the transfection was performed according to the manufacturer’s instructions. All downstream functional experiments were conducted 24 hours after transfection.

### Siglecs binding Assay

B16-F10 and B16-Ova cells that stably expressed ST8Sia6-GFP were screened for the generation of Siglec-E ligands using Recombinant Mouse Siglec E-Fc Chimera (catalog no. 551506, BioLegend). The cells were plated in 12-well plates at a density of 100,000 cells per well. After ST8Sia6 plasmid transfection, they were first blocked with rat serum (catalog no. 10710C, Invitrogen) for 10 minutes on ice, then stained with recombinant Siglec-E for 30 minutes. After washing with PBS (catalog no. 21-030-CV, Corning), the cells were stained with APC anti-Siglec-E antibody (catalog no. 677106, BioLegend), PNA- Cy3 for 30 minutes at 4°C. Following another wash with FACS buffer, the cells were analyzed using the Attune NxT flow cytometer (Life Technologies). The frequency of Siglec ligand-expressing cells was determined using FlowJo-10.2 software. This process also repeated in porcine primary kidney and aorta endothelial cell lines after transfecting with both control and ST8Sia6 plasmid. For porcine kidney and aorta endothelial cell lines, we also screened for Siglec-9 and Siglec-10 ligands using recombinant human Siglec-9 and Siglec-10 Fc Chimeras (Catalog nos. 1139SL050 and 2130SL050, BD Pharmaceuticals). Cells, transfected with the ST8Sia6 plasmid, were plated at 100,000 cells per well in 12-well plates. After blocking for 10 minutes on ice, they were stained with recombinant Siglec-9 and Siglec-10 for 30 minutes. Following washes with PBS (Catalog no. 21-030-CV, Corning), cells were further stained with PE anti-human Siglec-10 and Pacific Blue™ anti-human Siglec-9 antibodies (Catalog nos. 347603 and 351507, BioLegend) for 30 minutes at 4°C and further wash by FACS buffer. Analysis was performed using the Attune NxT flow cytometer (Life Technologies) and FlowJo-10.2 software.

### Western blot analysis

Cells from different assay were lysed in RIPA Lysis Buffer (Thermo Scientific™, catalog no. 89900) on ice for 1 hour, with vortexing every 15 minutes. Protein concentration was measured using the Pierce™ BCA Protein Assay Kit (catalog no. 23225). Proteins were separated by 4– 20% SDS-PAGE, transferred to an Immobulin-P PVDF membrane (Millipore) using BioRad Wet/Tank Blotting Systems for 100 minutes at 100 volts, and blocked with 5% skim milk in Tris-buffered saline with 0.5% Tween-20 (pH 7.4, catalog no. IBB-171, Boston Bioproducts Inc). Membranes were probed with primary antibodies at 1:1000 and HRP-conjugated secondary antibodies at 1:5000 dilution. Detection was carried out with the ECL Western Blotting Detection Kit (Thermo Scientific™, catalog no. 32209). ChemiDoc Imaging System were used for imaging of protein exprassion. Primary antibodies included ST8Sia6 Rabbit anti-Human pAb (Invitrogen™, catalog no. PA548789), Ovalbumin Rabbit/IgG pAb (Thermo Scientific, catalog no. PA5-97525), β-Actin (D6A8) Rabbit mAb (Cell Signaling, catalog no. 8457), and GAPDH (D16H11) XP® Rabbit mAb (Cell Signaling, catalog no. 5174S). Anti-rabbit IgG, HRP-linked antibody (Cell Signaling, catalog no. 7074S).

### OT-1 CD8+ T cells mediated cytotoxic killing assay

To assess the impact of ST8Sia6 overexpression on T cell-mediated cytotoxicity, we performed co-culture experiments with various cell lines, including B16-F10, B16-Ova. Different effector-to-target ratios were used in these assays to evaluate how ST8Sia6 overexpression affects the efficiency of T cell-mediated killing. Therefore, OT-1 CD8+ T cells were harvested from the spleens of C57BL/6-Tg(TcraTcrb)1100Mjb/J (Strain #:003831) mice and express a transgene encoding a TCR that specifically recognizes the SIINFEKL peptide bound to MHC-I H-2kb. The spleens were processed by mashing them through a 70-μm filter (catalog no. 352350, Corning) and then treated with red blood cell lysis buffer (catalog no. R 7757, Sigma Aldrich) for 2 minutes. The cells were washed twice with complete DMEM media. For the target cells, stable ST8Sia6- overexpressing B16-F10, B16-Ova cells were prepared at a concentration of 80 × 10^4^ live cells for each 24-well plate, and this preparation was done 24 hours before co-culture. The B16-F10 target cells were pulsed with SIINFEKL peptide (0.5 µg/mL) for 2 hours at 37°C in 5% CO2, followed by washing with complete DMEM media to remove any unbound peptide. The OT-1 CD8+ T cells were diluted in 500 µL of complete media and added to the B16-F10 target cells at various target-to-effector (E:T) ratios of 1:5, 1:2, and 1:1, with the target cell count remaining constant. The co-cultures were incubated for 24 hours at 37°C in 5% CO^2^ to assess T cell-mediated cytotoxicity. A similar procedure was followed for the ST8Sia6-overexpressing B16-Ova cell line without pulsed with SIINFEKL peptide to evaluate OT-1 CD8+ T cell-mediated cytotoxicity. After the 24-hour co-culture, the supernatants were collected for estimating cytokine productions and stored at −80 °C for cytokine analysis. The OT-1 CD8+ T cells were then collected from each treatment condition for analyzing surface marker activation, such as CD69 and LAMP-1, where OT-1 T cells were stained with APC-labeled anti-mouse CD45, PE-labeled anti-mouse CD8, FITC-labeled anti-mouse CD69, and BV421-labeled anti-mouse LAMP-1. Furthermore, attached target cells (such as B16-F10 and B16-Ova cells) from each treatment condition were scraped off, and relative killing efficiency was determined by GFP positive cells through flow cytometry. Samples were recorded on Attune NxT Flow Cytometer (Thermo Fisher Scientific, Waltham, MA, USA), and data were analyzed using FlowJo™ v10 Software (BD Biosciences).

### Human PBMC-mediated cytotoxicity against porcine endothelial cells

The cytotoxic activity of human immune cells was evaluated against porcine kidney and aorta cell lines overexpressing ST8Sia6. Peripheral blood mononuclear cells (PBMCs) were isolated from non-identified human whole blood using Ficoll density gradient separation according to standard procedures (Cytiva Ficoll-Paque™ PLUS, catalog no. 17144003). Freshly isolated PBMCs were resuspended in complete RPMI-1640 medium (Gibco, catalog no. 21870076), supplemented with 10% fetal bovine serum (HyClone, catalog no. SH30910.03), 1% penicillin/streptomycin solution (Gibco, catalog no. 15140-122), 1% L-glutamine (200 mmol/L) (Gibco, catalog no. 25030-081), 1% sodium pyruvate (Corning, catalog no. 25-000-CI), 1% HEPES (1 mol/L) (Gibco, catalog no. 15630-080), 1% MEM nonessential amino acids (Corning, catalog no. 25-025-CI), and 0.1% 2- mercaptoethanol. The ST8Sia6 overexpressed porcine kidney and aorta endothelial cells were prepared at a density of approximately 80 × 10^4 live cells/well for each 24-well plate, and this preparation was done 24 hours prior to co-culture. The target cells were incubated with human PBMCs at a 1:5 target-to-effector (T:E) ratio for 24 hours at 37°C in 5% CO2 to assess immune cell-mediated cytotoxicity. Following the 24-hour co-culture period, supernatants were collected and stored at −80°C for subsequent cytokine measurement. PBMCs from each treatment condition were then analyzed for frequency of T cells and NK cells including surface activation and inactivation markers expression, using flow cytometry. Additionally, attached target cells (porcine kidney and aorta endothelial cells) from each treatment condition were harvested, and relative killing efficiency was determined by measuring GFP-positive cells through flow cytometry.

### Cytokine measurement by ELISA

Cell culture supernatants were harvested and analyzed for cytokines by ELISA techniques. For mouse IFN-γ ELISA, 96 well high affinity flat bottom plate (Catalog no. 3361, Corning) were coated overnight at 4°C with 25 ng/ml monoclonal rat anti-murine IFN-γ (BD Bioscience, #551216) diluted in carbonate coating buffer pH 9.6. IFNγ ELISAs antibody pairs for mouse were purchased from BD Biosciences, performed according to manufacturer’s protocol. For Human-IFN-γ, Human- Perforin and Human-Granzyme-B ELISA were performed with commercially available kit (Biolegend catalog no 430104, Cat no. 439207, Invitrogen catalog no. BMS2306) respectively. In all assays, mean OD450nm readings obtained from triplicate wells. ELISAs were read on a SpectraMax iD5 microplate reader (Molecular Devices) and data analyzed using SoftMax Pro version 7.0.2 software (Molecular Devices).

### Coagulation Factor Measurement in Instant Blood-Mediated Inflammation

Whole blood was collected from four individual donors using surface-heparinized BD Vacutainer® blood collection tubes (Catalog no. 367874, BD) to prevent clotting. To ensure proper mixing of the anticoagulant with the blood, the tubes were gently inverted several times. ST8Sia6 and Control-GFP transfected porcine kidney and aorta endothelial cells were then mixed with 500 µL of whole blood (without additional anticoagulant) in triplicate conditions. The mixtures were incubated on a rocking device at 37°C for periods ranging from 0 minutes to 6 hours. Blood samples from each treatment condition were collected at 0, 3, and 6 hours. At each time point, blood was transferred into Eppendorf tubes, and plasma was isolated by centrifugation at 4000 rpm for 10 minutes at 4°C. The plasma samples were subsequently stored at -80°C until assayed. Plasma levels of thrombin/anti-thrombin III complex (TAT), C3a, C4d, and sC5b-9 were measured using commercially available ELISA kits: Human Thrombin-Antithrombin Complex ELISA Kit (Abcam AB1089071), Human C4b (Novus Biologicals™ NBP270046), Human C3a (Novus Biologicals™ NBP266755), and Human C5b-9 (Novus Biologicals™ NBP266708).

### Staining and flow cytometry analysis

PBMCs from the ST8Sia6 overexpression and Control-GFP experimental conditions were stained with various surface markers. To identify activation and inactivation surface markers on T cells and NK cells, the following commercially available antibodies were used, (all from Biolegend): PE/Dazzle 594 anti-Human CD94 (CloneDX22, 305519), PE/Cyanine7 Anti-Mouse/Human KLRG1 (Clone2F1/KLRG1, 138415),Brilliant Violet 510 Anti-Human CD28 (Clone CD28.2, 302935), Brilliant Violet 605 Anti-Human CD69 (Clone FN50, 310937), APC Anti-Human CD3 (Clone OKT3, 317317, Brilliant Violet 605 Anti-Human NKG2D (Clone 1D11, 320831),PE Anti-Human 2B4 (Clone C1.7, 329507), Brilliant Violet 510 Anti-Human NKp46 (Clone 9E2, 331923), Brilliant Violet 711 Anti-Human DNAM-1 (Clone 11A8, 338333), Brilliant Violet 421 Anti-Human CD122 (Clone TU27, 339009), PE/Cyanine7 Anti-Human CD160 (Clone BY55, 341211), PE Anti-Human CD4 (Clone SK3, 344605), Alexa Fluor 647 Anti-Human CD8 (Clone SK1, 344725), Pacific Blue Anti-Human CD25 (Clone M-A251, 356129), Alexa Fluor 488 Anti-Human CD56 (Clone 5.1H11, 362517), Brilliant Violet 711 Anti-Human PD-1 (Clone NAT105, 367427), Brilliant Violet 785 Anti-Human LAG-3 (Clone 11C3C65, 369321), PerCP/Cyanine5.5 Anti-Human CD107a (Clone H4A3, 328615), PE Anti-Human Siglec-10 (Clone 5G6, 347603), Pacific Blue Anti-Human Siglec-9 (Clone K8, 351507), PE Anti-Human CD56 (Clone 5.1H11, 362507), PE/Cyanine7 anti-human CD8 Antibody (Clone SK1, 344712), Brilliant Violet 421™ anti-human CD3 Antibody (Clone UCHT1, 300434), Brilliant Violet 605™ anti-human CD8a Antibody (Clone RPA-T8, 301040),Brilliant Violet 650™ anti-human CD19 (Clone HIB19, 302238), Brilliant Violet 711™ anti-human CD19 (Clone HIB19, 302246), Brilliant Violet 510™ anti-human CD4 (Clone OKT4, 317444), PE/Cyanine5 anti-human CD56 (Clone 5.1H11, 362516), FITC Anti-Human 2B4 (Clone 2-69, 393509), PE Anti-Human CD45 (Clone HI30, 982322). In all experiments, PBMCs were initially stained with eFluor780 fixable viability dye (catalog no. 65086514, Thermo Fisher Scientific) to exclude dead cells before surface staining. For surface staining, PBMCs from different experimental conditions were analyzed using two separate antibody panels for NK cells and T cells. For all spectral flow cytometry experiments, single stain compensations were created using UltraComp eBeads Plus Compensation Beads (Invitrogen) or heat-killed cell samples. Samples were analyzed on an Attune NxT Flow Cytometer (Thermo Fisher Scientific, Waltham, MA, USA), and the data were processed using FlowJo™ v10 Software (BD Biosciences).

### Estimating ITAM and ITIM expression in PBMCs Co-Cultured with ST8Sia6 Porcine Cells

The expression of ITAM and ITIM receptors in protein level, peripheral blood mononuclear cells (PBMCs) was analyzed after co-culturing with ST8Sia6-expressing porcine kidney and aorta cells, described in above methods section. This investigation aimed to understand the immunomodulatory effects of the sialyltransferase ST8Sia6 on human immune cells. Following the 24-hour co-culture period, PBMCs from each treatment condition from four individual donor with average five to six replica were then analyzed for ITAM and ITIM expression on T cells, NK cells and B cells, including helper T cells and cytotoxic T cells, by surface and intercellular staining using flow cytometry. The following intra cellular antibodies were used: APC anti-human TIGIT (clone A15153G, catalog no. 372706) and PE anti-TREM-2 (clone 6E9, catalog no. 824806) to identify various immunoreceptor ITIM and ITAM domains on T cells, NK cells, and B cells. After surface staining, PBMCs were fixed and permeabilized using the eBioscience™ Foxp3 / Transcription Factor Staining Buffer Kit (catalog no. 00-5523-00). For staining controls, PE Rat IgG2b, κ Isotype Control (clone RTK4530, catalog no. 400607) and APC Mouse IgG2a, κ Isotype Control (clone MOPC-173, catalog no. 400221) were used. Additionally, the protein levels of ITAM and ITIM receptors in each treatment condition were evaluated using Western blot analysis. Where membranes were probed with anti-TREM2 (Cat no. NC2316447), anti-SHP-2 (Cat no. AF1894), anti-Phospho-SHP2 (AF3790), anti DAP12 (NC2233698), anti-TIGIT (501145264), anti SHP-1 (AF1878) primary antibody according to recommended dilutions. Fluorescent tag secondary antibodies were used for detection (IRdye 800cw donkey anti-gt, Cat no. NC9604570, IRdye 680rd goat anti-rabbit, Cat no. NC0252291).

### RNA Extraction, Library Preparation, and Sequencing

Porcine kidney endothelial cells overexpressing ST8Sia6 and control-GFP cells were collected from four individual experiment after 24 hours transfection for total RNA extraction. Similarly, cells overexpressing ST8Sia6 and control-GFP were collected from co-culturing with four individuals PBMCs as previously described. Cells from different experimental conditions were collected keeping in ice, maintaining temperature at 4°C, washed with cold PBS, and stored as cell pellets at -80°C for total-RNA isolation. Total RNA extraction was performed using the RNeasy Plus Mini Kit (Qiagen, Cat No. 74136) following the manufacturer’s protocol. RNA purity and concentration were measured using the Thermo Scientific™ NanoDrop™ 2000/2000c Spectrophotometer. The extracted total-RNA was then used for quality control, library preparation by using paired end Illumina totalRNA, and sequencing at the Center for Medical Genomics of the Indiana University School of Medicine, Indianapolis. A total of 10 ng of purified RNA was converted into NGS cDNA libraries, and library quality was assessed using capillary electrophoresis. Based on insert quality and concentration, the libraries were pooled in equimolar ratios and sequenced on an Illumina NovaSeq X Plus sequencing instrument according to the manufacturer’s instructions, generating 100-bp paired-end reads. The sequence reads were mapped to the reference genome using STAR (Spliced Transcripts Alignment to a Reference) ^88^. RNA-seq data quality was assessed by determining the number of reads falling into different annotated regions (exonic, intronic, splice junctions, intergenic, promoter, UTR, etc.) of the reference genes using bamUtils ^89^.

### Differential gene expression analysis

Low quality mapped reads (including reads mapped to multiple positions) were excluded and feature Counts ^90^ was used to quantify the gene level expression. Differential gene expression analysis was performed with edgeR ^91^. In this workflow, the statistical methodology applied uses negative binomial generalized linear models with likelihood ratio tests. The leading fold change is calculated as the average (root-mean-square) of the largest absolute log-fold changes between each pair of samples. Ideally, samples would cluster well within the main condition/treatment of interest.

### Gene Set Enrichment Analysis (GSEA)

Differential gene expression analysis was conducted using RNA-seq data containing log2 fold change and adjusted p-values from samples of Porcine kidney endothelial cells overexpressing ST8Sia6 and control-GFP cells were collected from four individual experiment condition and the results were visualized using dot plots, enrichment plots, and heatmaps. Similarly, cells overexpressing ST8Sia6 and control-GFP were collected from co-culturing with four individuals PBMCs with four individual’s experimental setup as previously described. Volcano plots were generated in R using the ‘ggplot2’ package (v.3.5.1). Genes were plotted based on their log2 fold change (x-axis) and -log10 adjusted p-value (y-axis). Threshold lines were added to distinguish upregulated (orange), downregulated (blue), and non-significant (gray) genes, with log2 fold change thresholds set at ±1.5 and adjusted p-value threshold at 0.05. Significant genes were labeled using ‘geom_label’ to enhance visualization.

Gene set enrichment analysis (GSEA) ^92,93^ was conducted using the GSEA desktop application (v.4.3.3). The input dataset consisted of expression values containing log2 fold change and adjusted p-values derived from GFP control and ST86ia plasmid overexpression samples, and cells overexpressing ST8Sia6 and control-GFP were collected from co-culturing with PBMCs. The analysis was performed against predefined gene sets from the Kyoto Encyclopedia of Genes and Genomes (KEGG) database and Hallmark pathways from the Molecular Signatures Database (MSigDB). Gene sets were considered significantly enriched if they had a nominal p-value < 0.05. Visualization of significant pathways related to metabolic and immune cell components was achieved using dot plots, generated with the ‘ggplot2’ package in R (v.3.5.1), and heatmaps created using the Bioconductor package ‘ComplexHeatmap’ (v.2.22.0). For heatmap generation, expression values for GFP control and ST86ia plasmid overexpression samples, enabling visualization of pathway enrichment trends across experimental conditions to facilitate biological interpretation. To access interactive relationships, the identified DEGs from KEGG were mapped via STRING database (with the statistically significant interaction scores >0.5) and the Cytoscape software (version 3.10.3) was used to construct the protein-protein interaction networks between the DEGs.

### Statistical analysis

Statistical analysis was performed using GraphPad Prism 9.0. For comparison of two groups, two-tailed unpaired Student’s t tests or paired Student’s t tests were performed in different excrement. For comparison of more than two groups, one-way ANOVA was performed. For Kaplan-Meier survival curves, the p values were calculated using the log-rank test. Data are shown as mean ± SEM. The correlation was analyzed using a Pearson correlation test. P < 0.05 were considered significant, and statistical significance was denoted with exact p value, not significant (p > 0.05). Each experiment was repeated independently at least three times and with multiple technical replicas.

### Data availability

The datasets generated during and/or analyzed during the current study are available from the corresponding author on reasonable request. The scRNA sequencing data that support the findings of this study have been deposited in the Gene Expression Omnibus (GEO) with the accession code XXXX. Authors declare that all other data are available within the manuscript and its supplementary information files. Source data are provided with this paper. No custom code or mathematical algorithm is deemed central to the conclusions of this paper.

## Acknowledgements

The authors thank Dr. Amanda Poholek and their lab members at UPMC Children’s Hospital, University of Pittsburgh, Pennsylvania, for providing the B16-F10 and B16-Ova cell lines. Authors also thank to Dr. Greg M. Delgoffe, and their lab members at Department of Immunology, University of Pittsburgh, Pennsylvania for providing spleen samples from C57BL/6-Tg (TcraTcrb)1100Mjb/J mice for our research. We also extend our gratitude to the IU Genomics Core Facility and Bioinformatics Center for their assistance with bulk RNA sequencing and preliminary data analysis. This study was funded by the National Institute of Diabetes and Digestive and Kidney Diseases (NIDDK) under project 7R01DK132583. Additionally, the authors appreciate the support of the Center for Diabetes and Metabolic Diseases (CDMD) for providing access to its instrumental facilities.

## Contributions

S.R., J.D.P., R.B., and D.K.C.C. conceptualized the study. S.R. and J.D.P. designed all experimental studies. S.R. performing all the experiments and analyzing the data. S.R. and J.D.P. assisted in the planning of BulkRNA-seq experiments. C.R. contributed to the analysis of RNA sequencing data and generated all heatmaps through bioinformatics analysis. E.K.S., and D.K.C.C. participated in experimental discussions and study design. S.R. and J.D.P. wrote the first draft of manuscript, while S.R and C.R. wrote RNS-seq experimental part. E.K.S., D.K.C.C., R.B., and J.D.P. contributed to manuscript editing and finalization. All the authors contributed to the preparation of the manuscript.

---

**Supplementary Figure 1.**
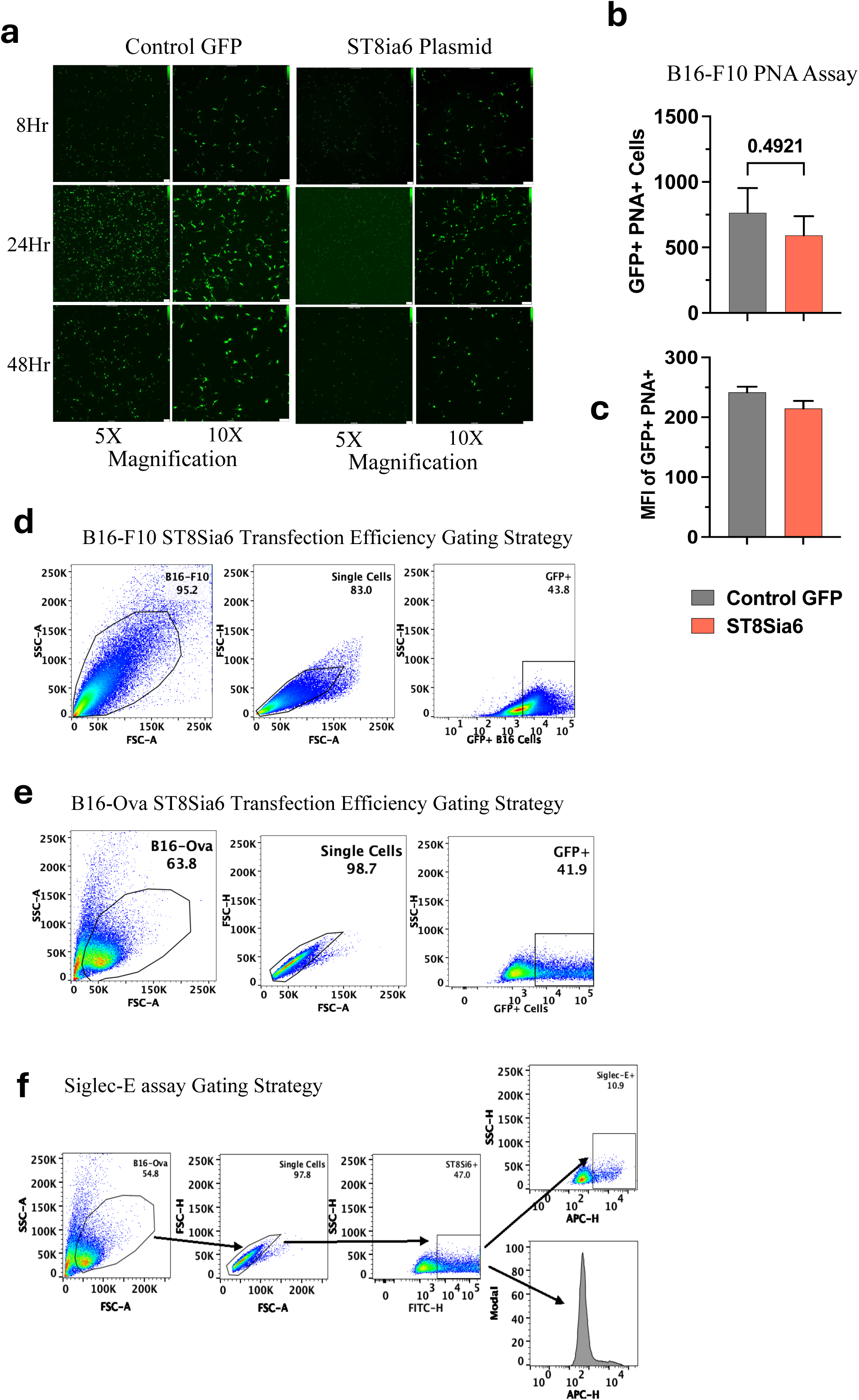
**a**, GFP fluorescent images at different time point after overexpression of ST8Sia6 in B16-F10 cell line**. b-c,** Flowcytometric analysis of Peanut Agglutinin (PNA) lectin positive cells number and MFI of ST8Sia6 plasmid or Control GFP transfect B16-F10 cells. **d-e,** Flow cytometry gating strategy for analysis of ST8Sia6 transfection efficiency in B16-F10 and B16-Ova cells. **f,** Flow cytometry gating strategy to assess the Siglec-E binding efficiency of ST8Sia6 transfected B16-F10 and B16-Ova cells. Significance analyzed by two-tailed unpaired t test between groups, significant different (P<0.05).

**Supplementary Figure 2.**
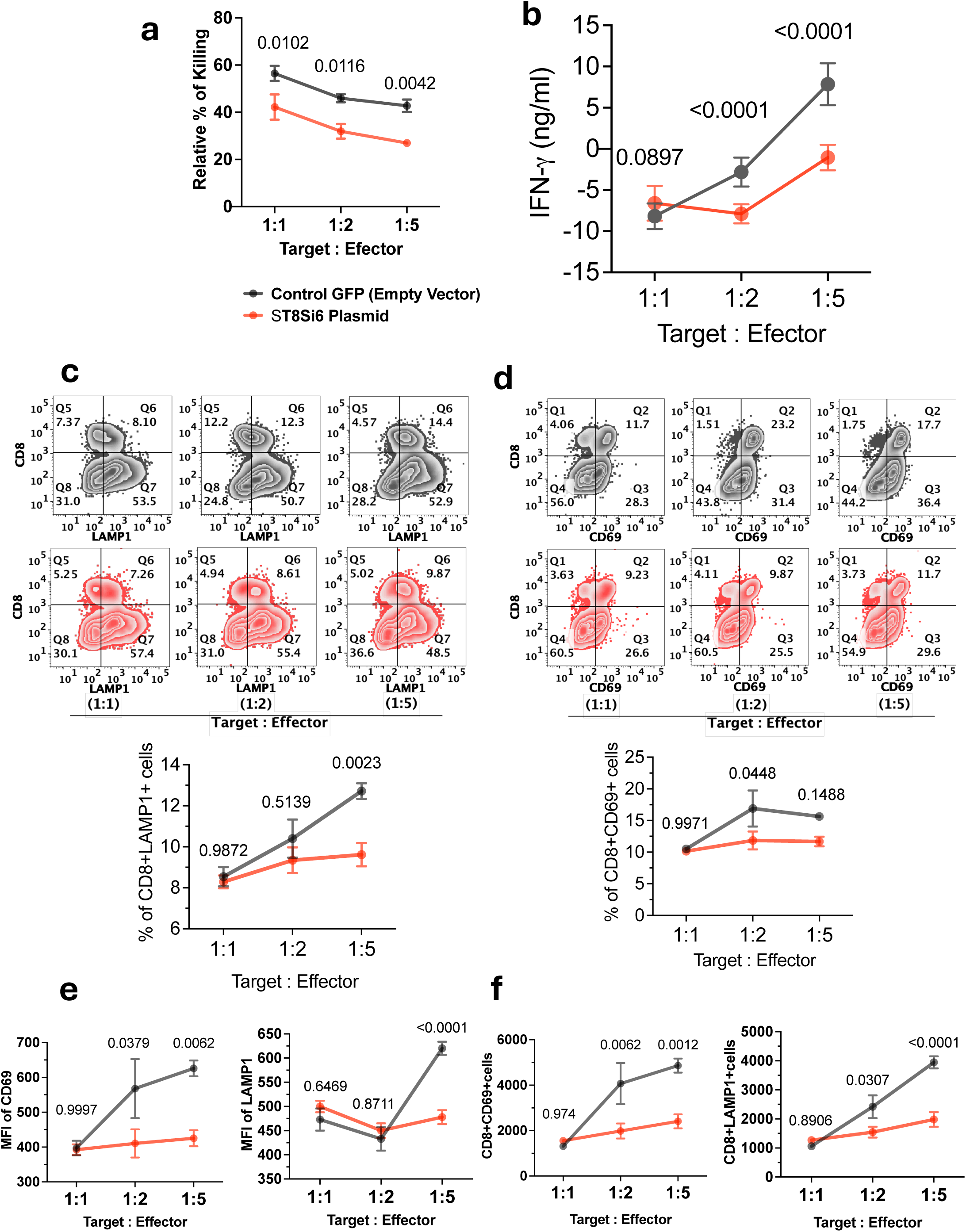
Sialic acid-overexpressing B16-Ova cells exhibit reduced cytotoxic effects mediated by OT-1 CD8+ T cells. In experimental conditions, B16-Ova cells transfected with a control GFP plasmid (grey curve) and those transfected with ST8Sia6 plasmid (red curve) were cocultured with OT-1 CD8+ T cells, which are specific effector cells, at various effector to target (E:T) ratios. The cytotoxicity was assessed relative to maximal cytotoxic effects. **a,** The percentage of killing efficiency of viable GFP-expressing target cells indicate the extent of effector cell-mediated cytotoxicity across different (E:T) ratios. **b,** IFN-γ levels were evaluated using ELISA from the supernatant of B16-Ova cocultures at different (E:T) ratios. **c-d,** Flowcytometry counter plot represent the percentage of CD8+CD69+ cells and CD8+LAMP1+ cells population respectively at various effector to target (E:T) ratios conditions. **e,** MFI of CD69 and LAMP1 expression in CD8+T cells across different E:T conditions. **f,** Absolute number of CD8+CD69+ CD8+LAMP1+ expressing cells across different E:T conditions. Data is representative of three independent experiments (mean ± SEM). E: T ratio = effector: target ratio; MFI = Mean Fluorescence Intensity. Significant difference assessed by two-way ANOVA and multiple comparison test between groups, significant different (P<0.05).

**Supplementary Fig 2.**
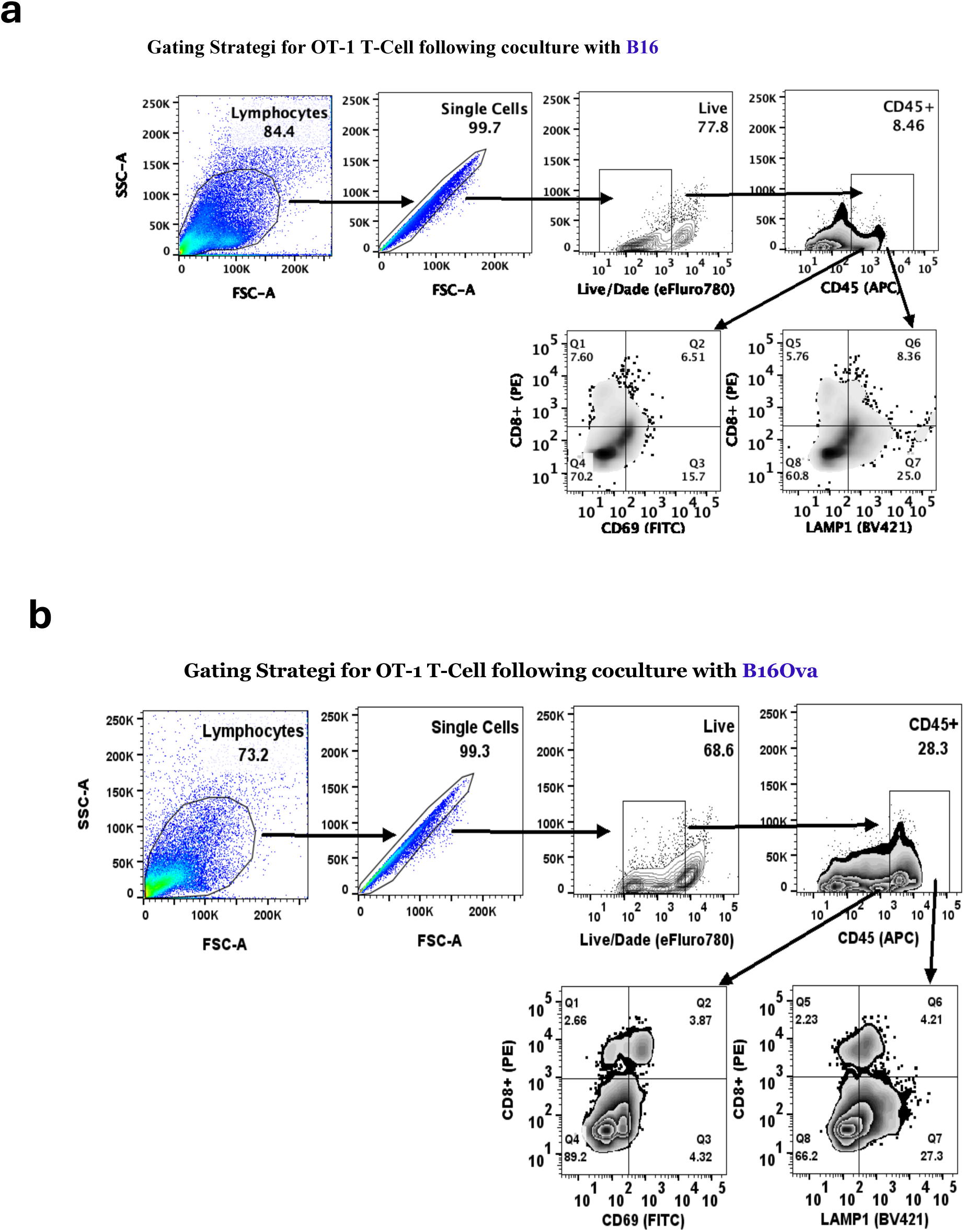
Overexpression of ST8Sia6 in B16-F10 and B16-Ova cells suppress OT-1 CD8+ T cells mediated cytotoxic effects. Gating strategy used in flow cytometry analysis to measure different activation marker for OT-1 CD8+ T cells. Splenocyte populations were gated on a forward scatter (FSC)/side scatter (SSC) plot. Splenocytes were then further gated on Live cells and CD45+ cells to determine CD8+CD69+ and CD8+LAMP1+ cell populations. Sample from B16-F10 **(e)** and B16-Ova **(f)** are shown in these diagrams.

**Supplementary Figure 4.**
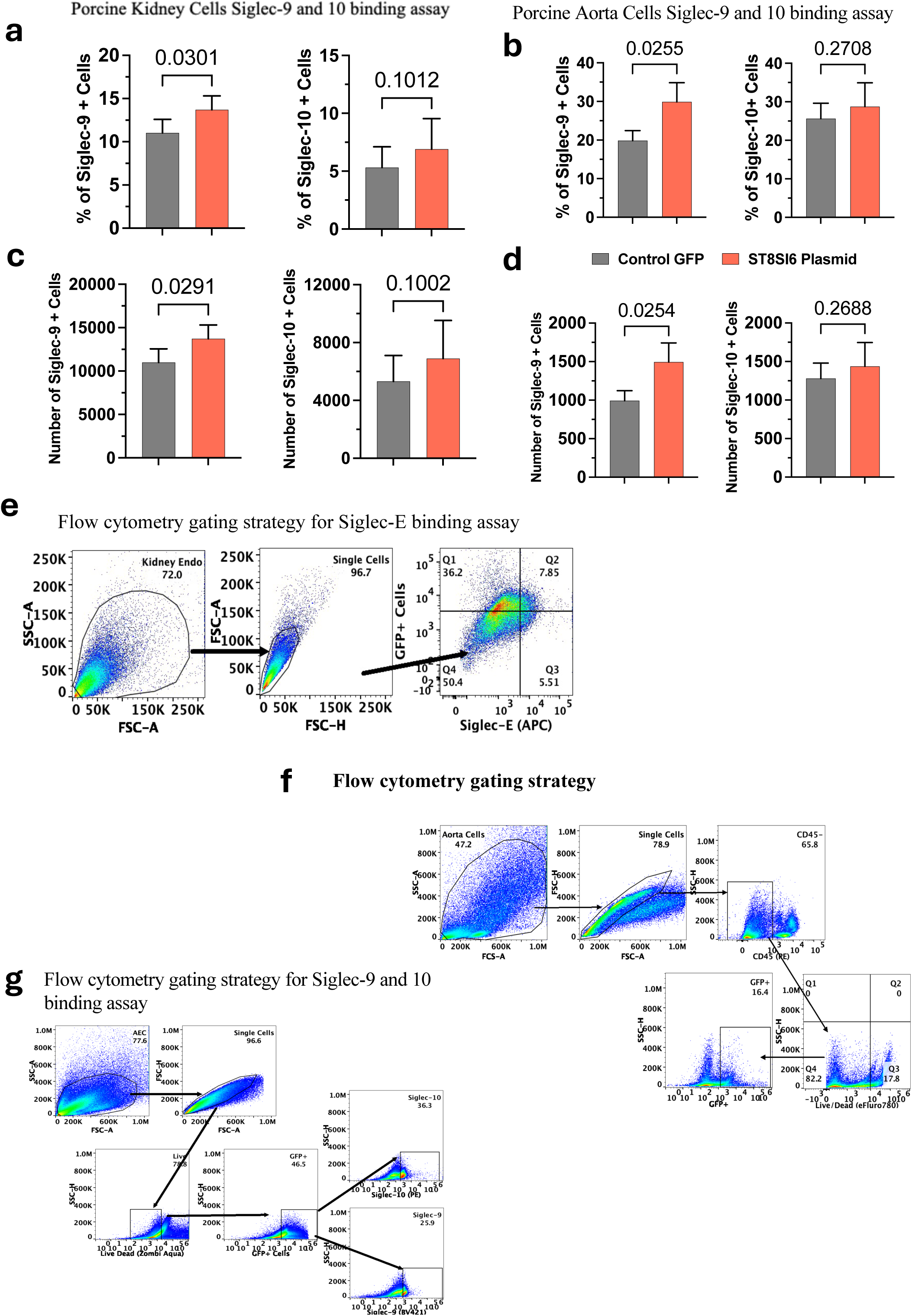
Siglec-9 and Siglec-10 binding assays were performed by probing for ligands using recombinant human Siglec-9, and Siglec-10, the expression levels were assessed via flow cytometry analysis with an anti-human Siglec-9 and anti-human Siglec- 10 antibody. a-b, Represented percentage of Siglec-9 and Sigle-10 ligands positive porcine kidney and aorta cells respectively. c-d, Absolute number of Siglec-9 and Siglec-10 positive porcine kidney and aortic endothelial cells compared with control GFP transfected experimental group by using flow cytometry. Data are presented as the mean ± SE from four independent experiments. Significant difference assessed by unpaired t-test. e. Flowcytometry plot represent gating strategy of Siglec-E binding assay. f, Gating strategy for estimating transfection efficiency. g, Flowcytometry plot represent gating strategy of Siglec-9 and Siglec-10 ligand binding assay analysis.

**Supplementary Figure 5.**
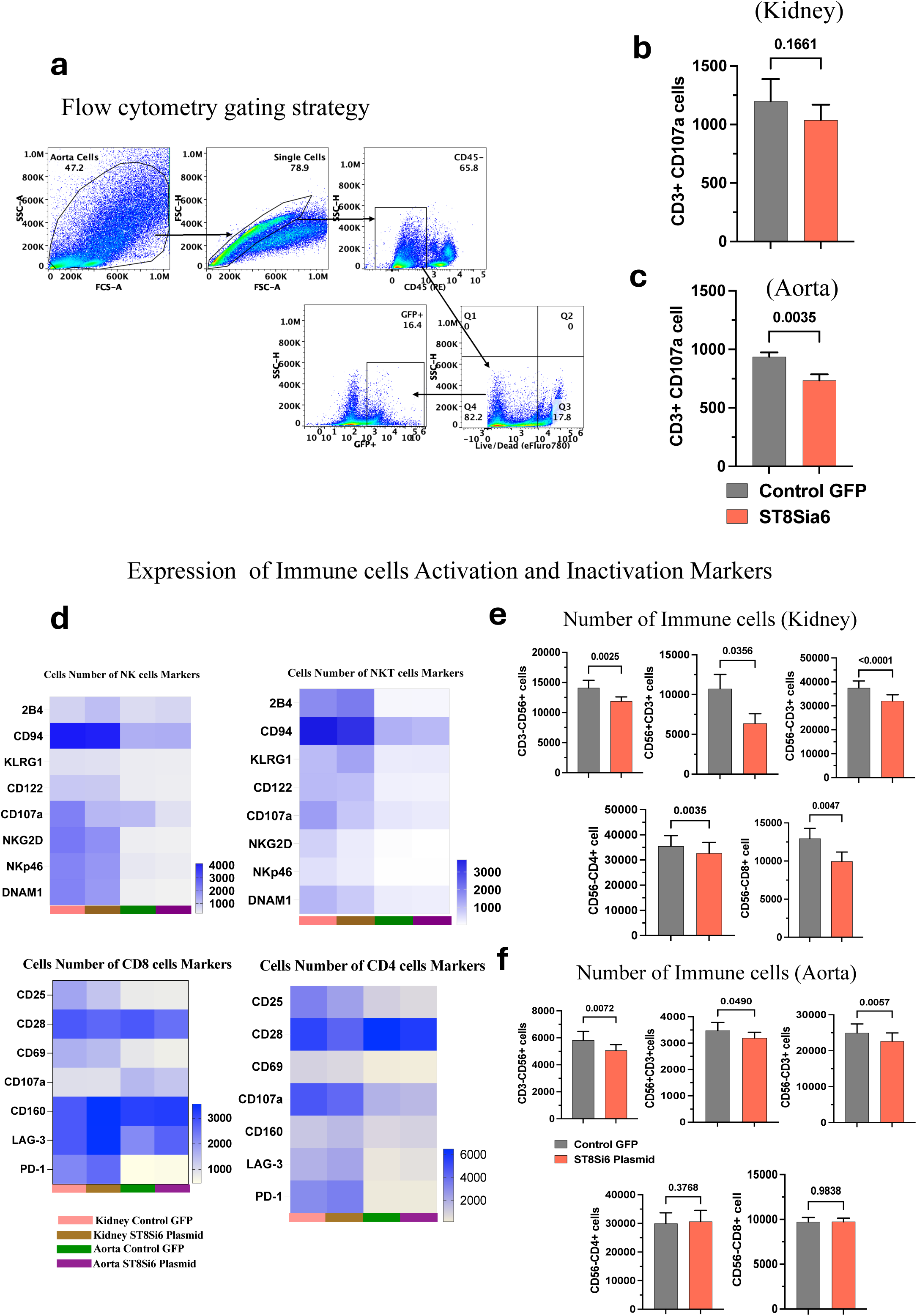
**a**, Flowcytometry plot represent gating strategy of porcine endothelial (Kidney / Aorta) from PBMC co-culture experiments, where porcine cells gated on a forward scatter (FSC)/side scatter (SSC) plot, single cell population, CD45- cells and Live cells population. Live cells were then further gated on to determine GFP+ cell populations. **b-c,** Absolute Number of CD3+ CD107a+ cell in different treatment condition. **d,** Hitmap represent absolute numbers of different activation and inactivation surface marker expressing cells in NK cell and T cells subpopulations. **e-f,** Compression of absolute number of different immune cells populations from ST8Sia6 transfected porcine kidney and aorta coculture conditions. Graphs show data from 4 donors (mean ± S.E) with 4 replicates. Significant differences (P<0.05) were determined by paired t-test, and P values are displayed in the graphs.

**Supplementary Figure 6.**
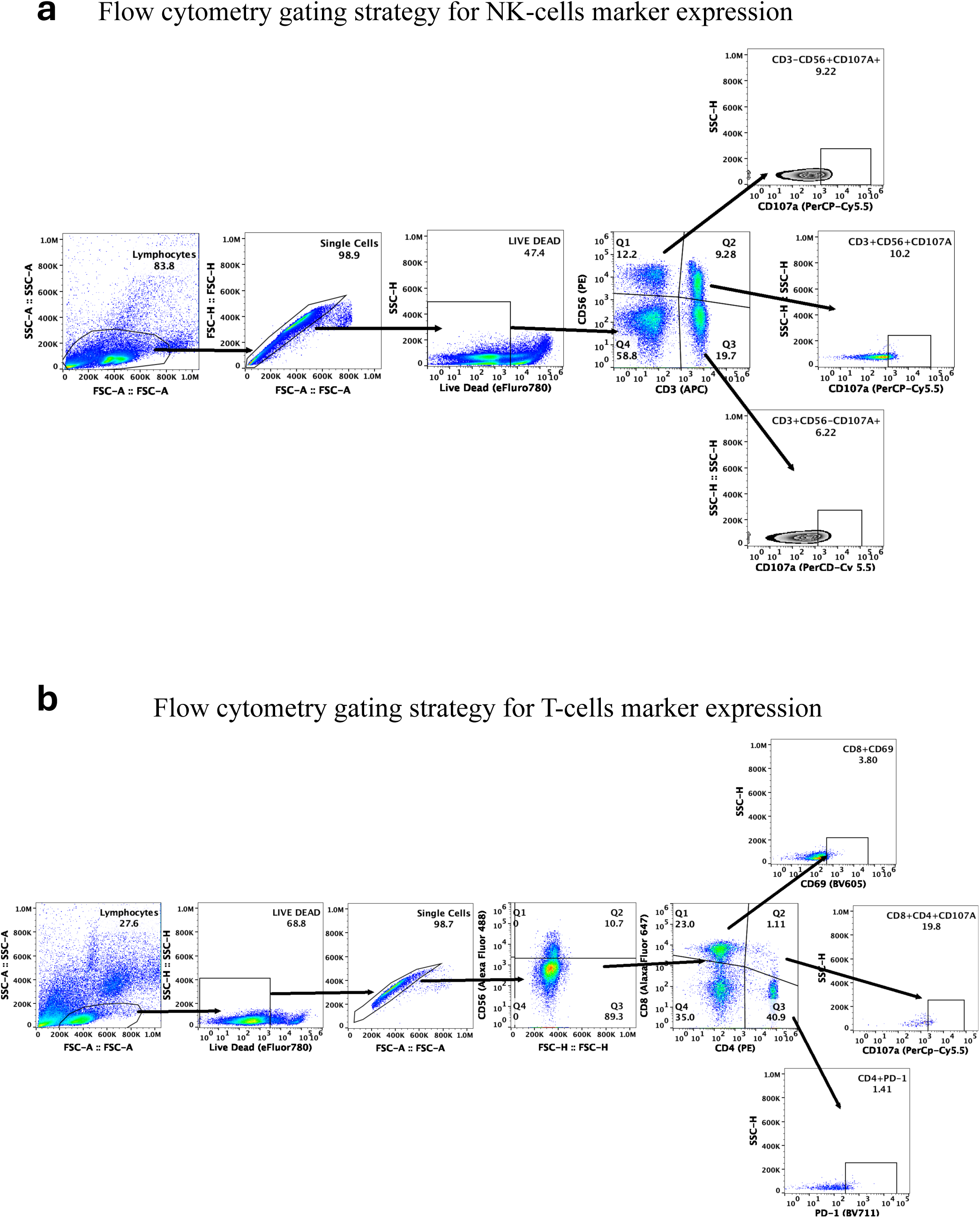
Flowcytometry plot represent gating strategy of PBMC from porcine endothelial (Kidney / Aorta) and PBMC co-culture experiments. a, PBMC cells gated on a forward scatter (FSC)/side scatter (SSC) plot, single cell population, live cells population. Live cells were then further gated on to CD56+ (NK) and CD3 (T cell) cell populations, further gated from CD56+ (NK), CD56+CD3+ (NKT) and CD3+ (T cells) to estimate the different activation and inactivation marker expressing subpopulations. **b,** PBMC cells gated on a forward scatter (FSC)/side scatter (SSC) plot, live cells population and then open for single cell population. Single cells were then further gated on to CD56- (NK) and open for CD4+ and CD8+ (T cell) cell populations, after that gated further on CD4+ T-cells, CD8+ T cells and CD4+CD8+ T cells to estimate the different activation and inactivation marker expressing subpopulations.

**Supplementary Figure 7.**
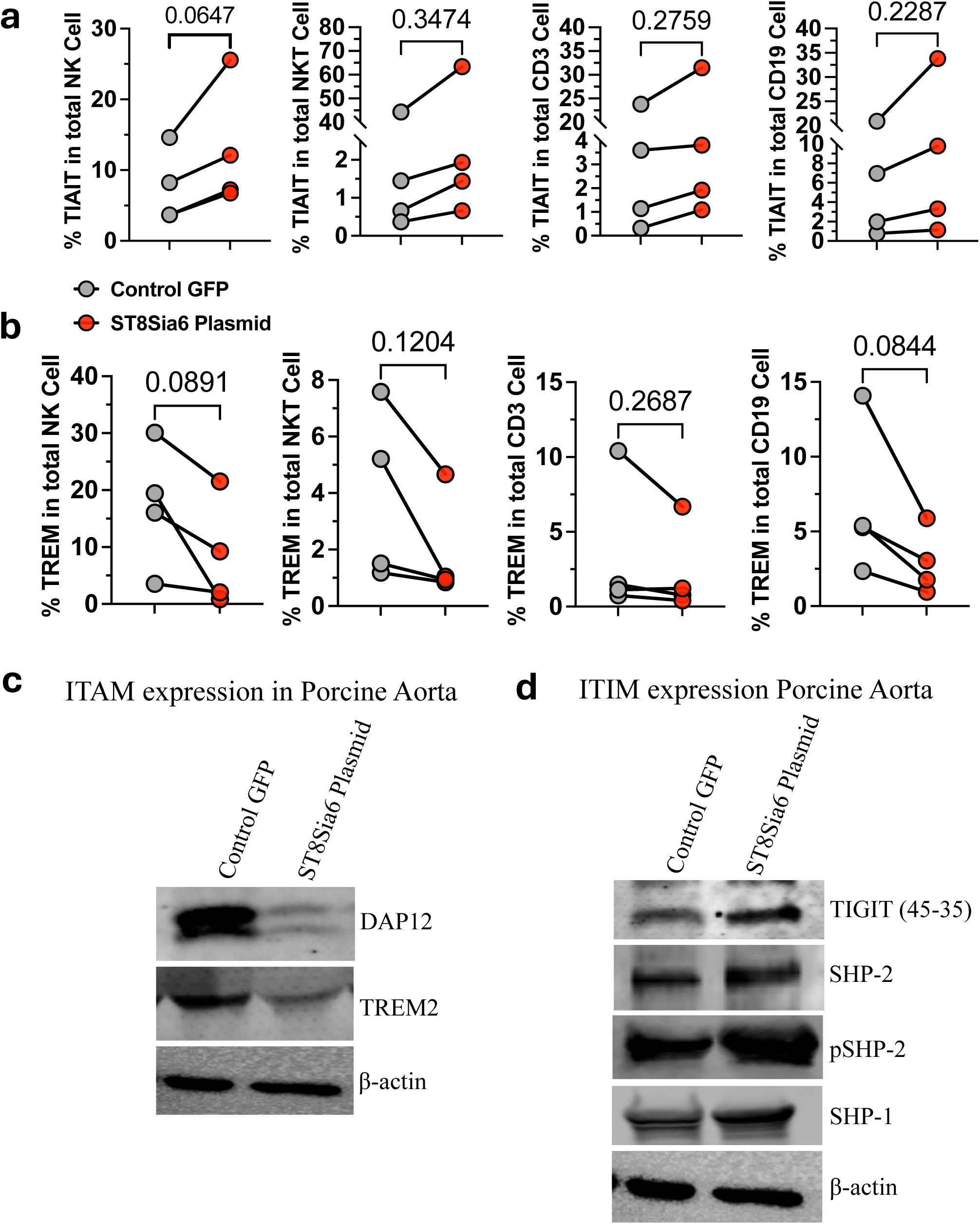
The overexpression of sialic acid in porcine aorta endothelial cells primarily regulate immune responses through ITAM/ITIM signaling pathways in various immune cells. a, The percentage of a tyrosine-based inactivation motif TIAIT expression in different immune cells within the ST8Sia6 and Control GFP-expressing groups. b, The proportion of a tyrosine-based activation motif TREM expression across various immune cell types in the ST8Sia6 group compared to the Control GFP-expressing group. c-d, Evaluation of ITAM and ITIM protein expression in PBMC lysates from the aorta co-culture experiment probe with anti-DAP12, anti-TREM2 primary antibody for detection of ITAM and anti TIGIT, anti-SHP-2, anti pSHP-2, anti- SHP-1, primary antibody for ITIM detection. Graphs show data from 4 donors (mean ± S.E) in average replicates. Significant differences (P<0.05) were determined by paired t-test, and P values are displayed in the graphs.

**Supplementary Figure 8.**
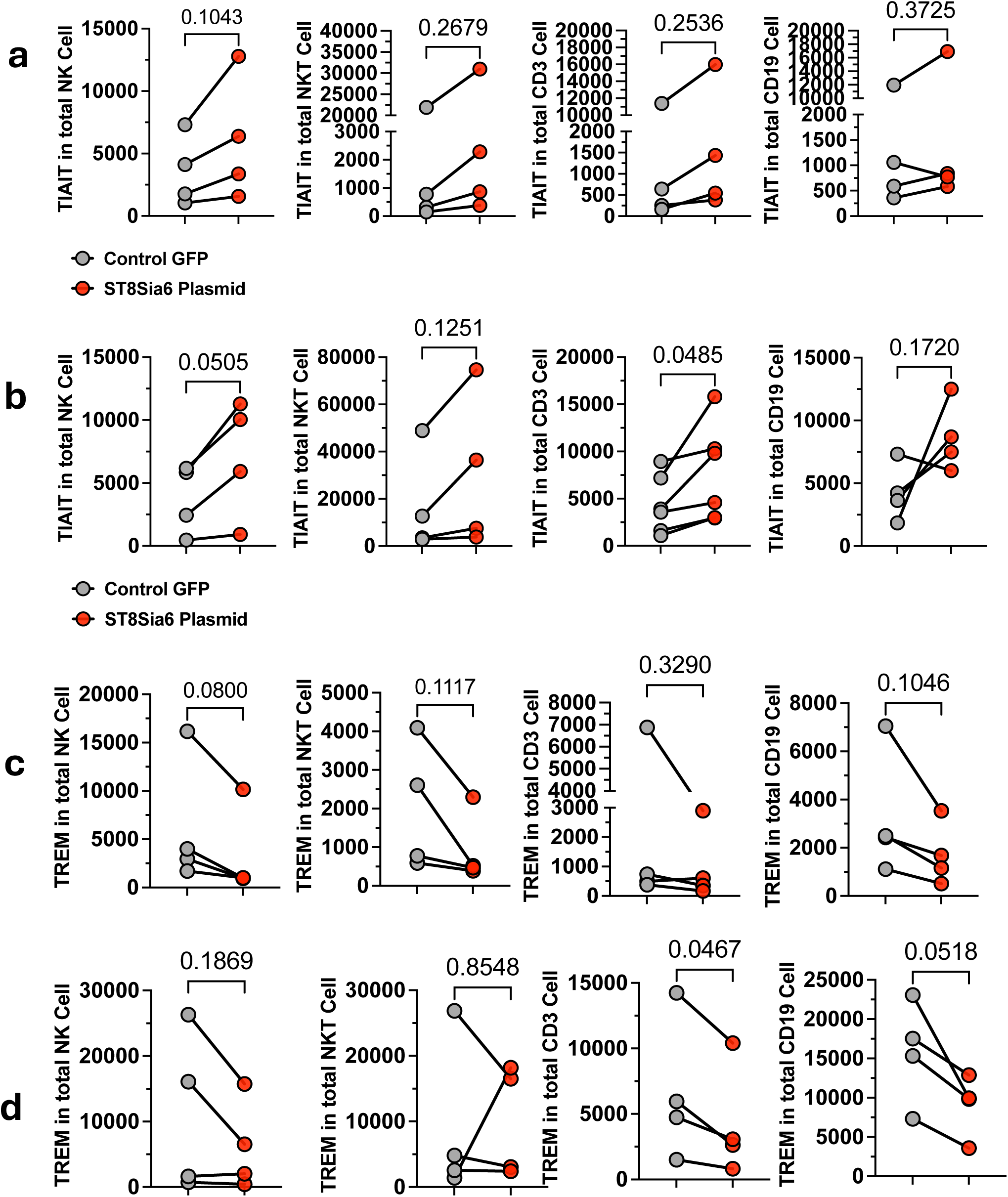
Sialic acid expression in porcine kidney and aorta endothelial cells modulates number of ITAM and ITIM expressing cells across different immune cell types. a-b, Absolute number of TIAIT expressing immune cells in aorta and kidney coculture conditions. c-d, Absolute number of TREM expressing different immune cells from aorta and kidney coculture conditions. Graphs show data from 4 donors (mean ± S.E) in average replicates. Significant differences (P<0.05) were determined by paired t-test, and P values are displayed in the graphs.

**Supplementary Figure 9.**
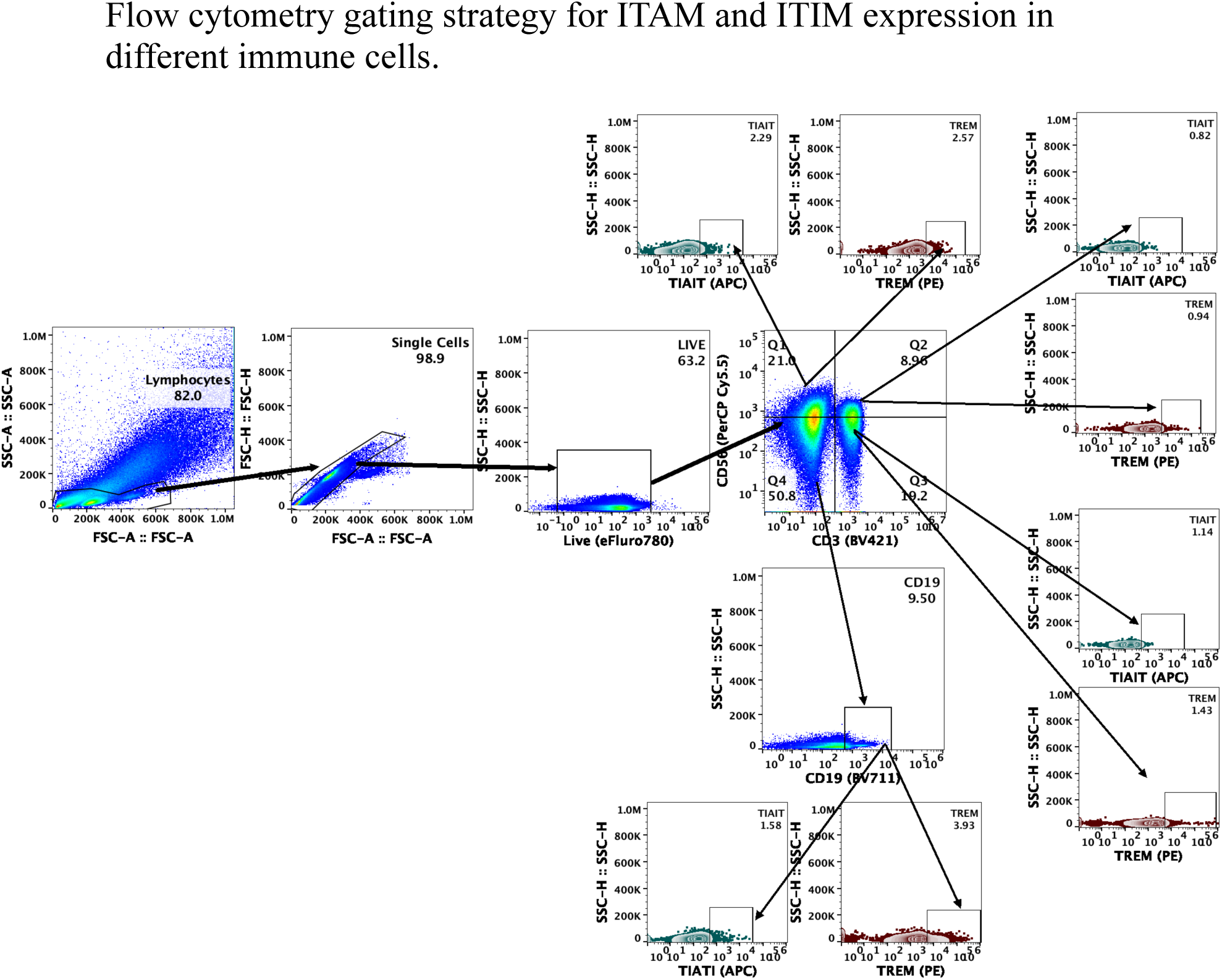
Flowcytometry plot represent gating strategy for ITAM and ITIM expression in immune cell population from human PBMC after co-culture with ST8Sia6 transfected porcine endothelial (Kidney / Aorta) cells. PBMC cells gated on a forward scatter (FSC)/side scatter (SSC) plot, single cell population, live cells population. Live cells were then further gated on to CD56+ (NK) and CD3 (T cell) cell populations, further gated from CD56+ (NK), CD56+CD3+ (NKT), CD3+ (T cells) and CD19 (B cell) to estimate the different Immunoreceptor Tyrosine-based Activation Motif (ITAM) and Immunoreceptor Tyrosine-based Inactivation Motif (ITIM) marker expression.

**Supplementary Figure 10.**
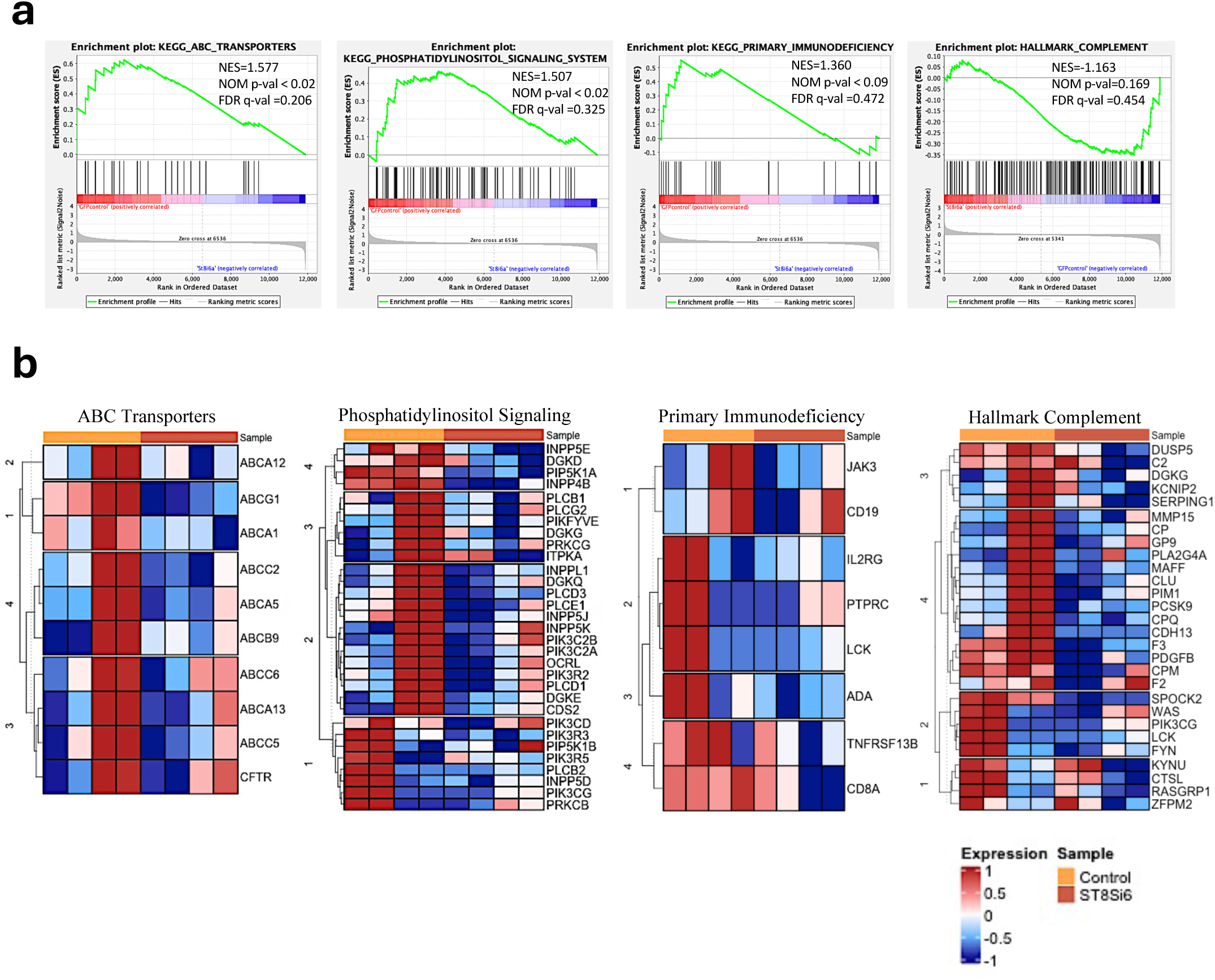
**a.** GSEA enrichment plots comparing ST8Sia6 overexpression vs GFP control. Pathways analyzed include ABC Transporters, phosphatidylinositol signaling, primary Immunodeficiency, and Complement genes. **b.** Heat map of the top 30 marker genes for each pathway in the comparison of ControlGFP (Yellow, left three column) vs. ST8Sia6 overexpress (Orange, right three column). Red, upregulated; Blue, downregulated genes.

**Supplementary Figure 11.**
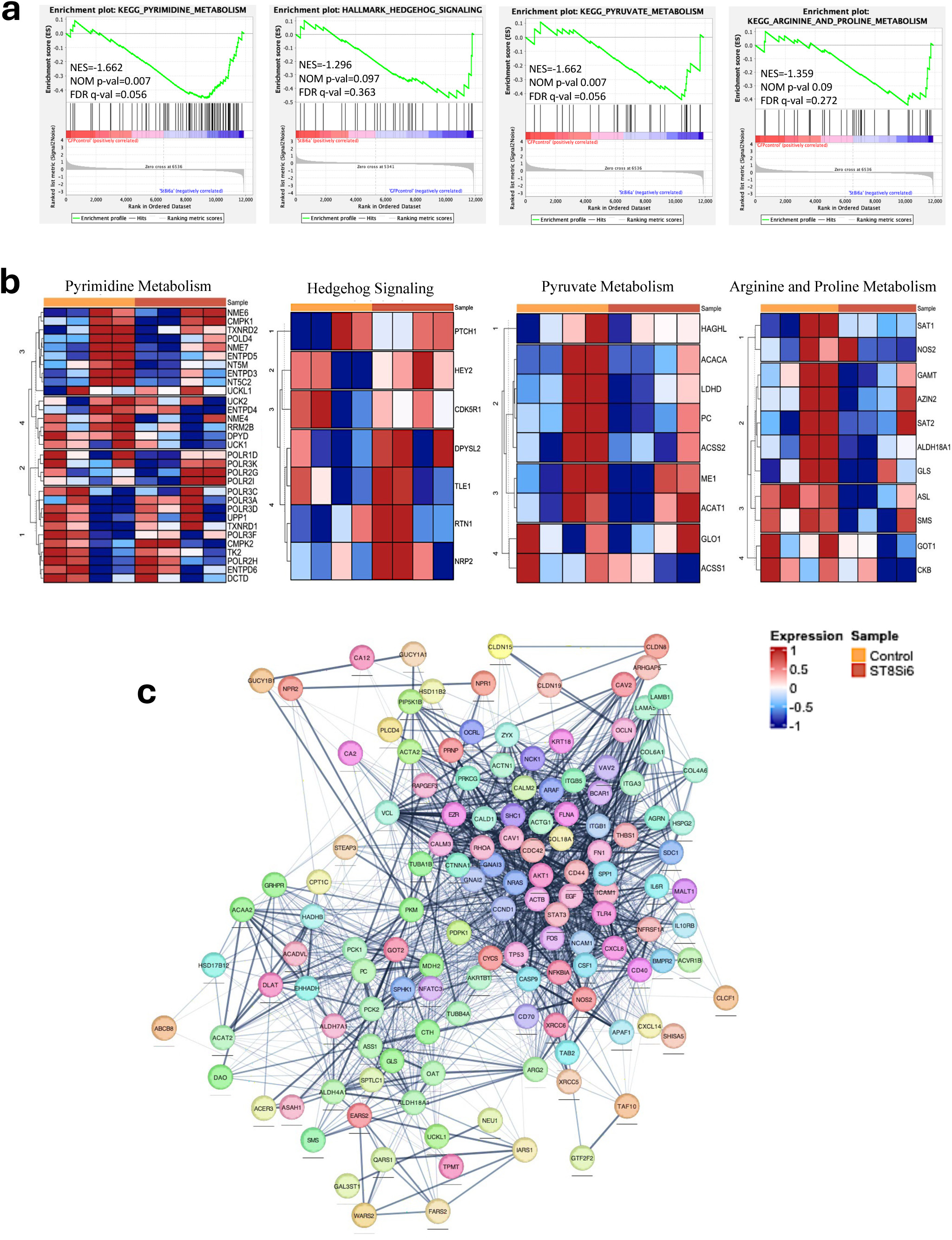
**a-b.** GSEA analysis of pathways involved in Pyrimidine Metabolism, Hedgehog Signaling, Pyruvate Metabolism, and Arginine and Proline Metabolism. The graphs show the NES and FDR, while the heatmaps highlight the genes that are differentially expressed within each gene set. **c**. Cytoscape enrichment map of over- and under-expressed genes in the enriched GSEA pathways in ST8Sia6 overexpression compared to GFP control. Each dot represents a gene enriched in the pathway, and the lines represent interconnectedness. Groupings of the genes in the pathways are labeled by common functions.

## References

1. Carrier, A.N., et al. Xenotransplantation: A New Era. Front Immunol 13, 900594 (2022).

2. Gonzalez, J.M., et al. Evaluating the Performance of a Nonelectronic, Versatile Oxygenating Perfusion System across Viscosities Representative of Clinical Perfusion Solutions Used for Organ Preservation. Bioengineering (Basel) 10(2022).

3. Shelton, B.A., et al. Increasing Obesity Prevalence in the United States End-Stage Renal Disease Population. J Health Sci Educ 2(2018).

4. Bezinover, D. & Saner, F. Organ transplantation in the modern era. BMC Anesthesiol 19, 32 (2019).

5. Reardon, S. First pig-to-human heart transplant: what can scientists learn? Nature 601, 305–306 (2022).

6. Siems, C., Huddleston, S. & John, R. A Brief History of Xenotransplantation. Ann Thorac Surg 113, 706–710 (2022).

7. David K.C. Cooper, L.W., Kohei Kinoshita, Zahra Habibabady, Ivy Rosales, Takaaki Kobayashi, Hidetaka Hara. Immunobiological barriers to pig organ xenotransplantation Authors. European Journal of Transplantation 1, 167–181 (2023).

8. Xi, J., et al. Genetically engineered pigs for xenotransplantation: Hopes and challenges. Front Cell Dev Biol 10, 1093534 (2022).

9. Cooper, D.K., Koren, E. & Oriol, R. Genetically engineered pigs. Lancet 342, 682–683 (1993).

10. Cowan, P.J. & Tector, A.J. The Resurgence of Xenotransplantation. Am J Transplant 17, 2531–2536 (2017).

11. Cooper, D.K.C., et al. Justification of specific genetic modifications in pigs for clinical organ xenotransplantation. Xenotransplantation 26, e12516 (2019).

12. Ma, D., et al. Kidney transplantation from triple-knockout pigs expressing multiple human proteins in cynomolgus macaques. Am J Transplant 22, 46–57 (2022).

13. Platts-Mills, T.A.E., et al. On the cause and consequences of IgE to galactose-alpha-1,3- galactose: A report from the National Institute of Allergy and Infectious Diseases Workshop on Understanding IgE-Mediated Mammalian Meat Allergy. J Allergy Clin Immunol 145, 1061–1071 (2020).

14. Adams, A.B., et al. Anti-C5 Antibody Tesidolumab Reduces Early Antibody-mediated Rejection and Prolongs Survival in Renal Xenotransplantation. Ann Surg 274, 473–480 (2021).

15. Cooper, D.K.C., Ekser, B. & Tector, A.J. Immunobiological barriers to xenotransplantation. Int J Surg 23, 211–216 (2015).

16. Vadori, M. & Cozzi, E. Immunological challenges and therapies in xenotransplantation. Cold Spring Harb Perspect Med 4, a015578 (2014).

17. Bach, F.H., et al. Modification of vascular responses in xenotransplantation: inflammation and apoptosis. Nat Med 3, 944–948 (1997).

18. Platt, J.L., et al. Immunopathology of hyperacute xenograft rejection in a swine-to-primate model. Transplantation 52, 214–220 (1991).

19. Rose, A.G., Cooper, D.K., Human, P.A., Reichenspurner, H. & Reichart, B. Histopathology of hyperacute rejection of the heart: experimental and clinical observations in allografts and xenografts. J Heart Lung Transplant 10, 223–234 (1991).

20. Cooper, D.K.C., Mou, L. & Bottino, R. A brief review of the current status of pig islet xenotransplantation. Front Immunol 15, 1366530 (2024).

21. Li, Y. & Chen, X. Sialic acid metabolism and sialyltransferases: natural functions and applications. Appl Microbiol Biotechnol 94, 887–905 (2012).

22. Varki, A. N-glycolylneuraminic acid deficiency in humans. Biochimie 83, 615–622 (2001).

23. Saini, P., Adeniji, O.S. & Abdel-Mohsen, M. Inhibitory Siglec-sialic acid interactions in balancing immunological activation and tolerance during viral infections. EBioMedicine 86, 104354 (2022).

24. Crocker, P.R., Paulson, J.C. & Varki, A. Siglecs and their roles in the immune system. Nat Rev Immunol 7, 255–266 (2007).

25. Abeln, M., et al. Sialic acid is a critical fetal defense against maternal complement attack. J Clin Invest 129, 422–436 (2019).

26. Hernandez-Arteaga, A.C., et al. Comparative study of sialic acid content in saliva between preeclampsia and normal gestation patients. Placenta 130, 12–16 (2022).

27. Egan, H., et al. Targeting stromal cell sialylation reverses T cell-mediated immunosuppression in the tumor microenvironment. Cell Rep 42, 112475 (2023).

28. van de Wall, S., Santegoets, K.C.M., van Houtum, E.J.H., Bull, C. & Adema, G.J. Sialoglycans and Siglecs Can Shape the Tumor Immune Microenvironment. Trends Immunol 41, 274–285 (2020).

29. Teintenier-Lelievre, M., et al. Molecular cloning and expression of a human hST8Sia VI (alpha2,8-sialyltransferase) responsible for the synthesis of the diSia motif on O- glycosylproteins. Biochem J 392, 665–674 (2005).

30. Friedman, D.J., et al. ST8Sia6 Promotes Tumor Growth in Mice by Inhibiting Immune Responses. Cancer Immunol Res 9, 952–966 (2021).

31. Zhu, W., Zhou, Y., Guo, L. & Feng, S. Biological function of sialic acid and sialylation in human health and disease. Cell Death Discov 10, 415 (2024).

32. Belmonte, P.J., Shapiro, M.J., Rajcula, M.J., McCue, S.A. & Shapiro, V.S. Cutting Edge: ST8Sia6-Generated alpha-2,8-Disialic Acids Mitigate Hyperglycemia in Multiple Low-Dose Streptozotocin-Induced Diabetes. J Immunol 204, 3071–3076 (2020).

33. Pillai, S., Netravali, I.A., Cariappa, A. & Mattoo, H. Siglecs and immune regulation. Annu Rev Immunol 30, 357–392 (2012).

34. Hsu, F.C., et al. Immature recent thymic emigrants are eliminated by complement. J Immunol 193, 6005–6015 (2014).

35. Zhong, L., et al. Hyperacute rejection-engineered oncolytic virus for interventional clinical trial in refractory cancer patients. Cell 188, 1119–1136 e1123 (2025).

36. Munkley, J. & Elliott, D.J. Hallmarks of glycosylation in cancer. Oncotarget 7, 35478–35489 (2016).

37. Dobie, C. & Skropeta, D. Insights into the role of sialylation in cancer progression and metastasis. Br J Cancer 124, 76–90 (2021).

38. Laubli, H., Nalle, S.C. & Maslyar, D. Targeting the Siglec-Sialic Acid Immune Axis in Cancer: Current and Future Approaches. Cancer Immunol Res 10, 1423–1432 (2022).

39. in Essentials of Glycobiology (eds. Varki, A., et al.) (Cold Spring Harbor (NY), 2022).

40. Muhlenhoff, M., Rollenhagen, M., Werneburg, S., Gerardy-Schahn, R. & Hildebrandt, H. Polysialic acid: versatile modification of NCAM, SynCAM 1 and neuropilin-2. Neurochem Res 38, 1134-1143 (2013).

41. Schnaar, R.L., Gerardy-Schahn, R. & Hildebrandt, H. Sialic acids in the brain: gangliosides and polysialic acid in nervous system development, stability, disease, and regeneration. Physiol Rev 94, 461–518 (2014).

42. David J Friedman, M.S., Matthew Rajcula, Shaylene McCue, Virginia Smith Shapiro. ST8Sia6 Overexpression Accelerates Tumor Growth, Alters Macrophage Polarization and the Immune Response. The Journal of Immunology 202, Supp.135.138 (2019).

43. Lin, C.H., Yeh, Y.C. & Yang, K.D. Functions and therapeutic targets of Siglec-mediated infections, inflammations and cancers. J Formos Med Assoc 120, 5–24 (2021).

44. Zhang, J.Q., Biedermann, B., Nitschke, L. & Crocker, P.R. The murine inhibitory receptor mSiglec-E is expressed broadly on cells of the innate immune system whereas mSiglec-F is restricted to eosinophils. Eur J Immunol 34, 1175–1184 (2004).

45. Eisenson, D.L., Hisadome, Y., Santillan, M.R. & Yamada, K. Progress in islet xenotransplantation: Immunologic barriers, advances in gene editing, and tolerance induction strategies for xenogeneic islets in pig-to-primate transplantation. Front Transplant 1(2022).

46. Naziruddin, B., et al. Evidence for instant blood-mediated inflammatory reaction in clinical autologous islet transplantation. Am J Transplant 14, 428–437 (2014).

47. Kourtzelis, I., Magnusson, P.U., Kotlabova, K., Lambris, J.D. & Chavakis, T. Regulation of Instant Blood Mediated Inflammatory Reaction (IBMIR) in Pancreatic Islet Xeno- Transplantation: Points for Therapeutic Interventions. Adv Exp Med Biol 865, 171–188 (2015).

48. Maeda, A., et al. The Innate Cellular Immune Response in Xenotransplantation. Front Immunol 13, 858604 (2022).

49. Cardone, J., et al. Complement regulator CD46 temporally regulates cytokine production by conventional and unconventional T cells. Nat Immunol 11, 862–871 (2010).

50. Varki, A. Sialic acids in human health and disease. Trends Mol Med 14, 351–360 (2008).

51. Pontrelli, P., et al. The Role of Natural Killer Cells in the Immune Response in Kidney Transplantation. Front Immunol 11, 1454 (2020).

52. Puga Yung, G., Schneider, M.K.J. & Seebach, J.D. The Role of NK Cells in Pig-to-Human Xenotransplantation. J Immunol Res 2017, 4627384 (2017).

53. Billadeau, D.D. & Leibson, P.J. ITAMs versus ITIMs: striking a balance during cell regulation. J Clin Invest 109, 161–168 (2002).

54. Laubli, H. & Varki, A. Sialic acid-binding immunoglobulin-like lectins (Siglecs) detect self-associated molecular patterns to regulate immune responses. Cell Mol Life Sci 77, 593–605 (2020).

55. Bender, C., et al. Islet-Expressed CXCL10 Promotes Autoimmune Destruction of Islet Isografts in Mice With Type 1 Diabetes. Diabetes 66, 113–126 (2017).

56. Ko, C.Y., et al. Bioinformatics Analyses Identify the Therapeutic Potential of ST8SIA6 for Colon Cancer. J Pers Med 12(2022).

57. Yuan, Y., et al. Complement networks in gene-edited pig xenotransplantation: enhancing transplant success and addressing organ shortage. J Transl Med 22, 324 (2024).

58. Ekser, B., Cooper, D.K.C. & Tector, A.J. The need for xenotransplantation as a source of organs and cells for clinical transplantation. Int J Surg 23, 199–204 (2015).

59. Ryczek, N., Hryhorowicz, M., Zeyland, J., Lipinski, D. & Slomski, R. CRISPR/Cas Technology in Pig-to-Human Xenotransplantation Research. Int J Mol Sci 22(2021).

60. Lutz, A.J., et al. Double knockout pigs deficient in N-glycolylneuraminic acid and galactose alpha-1,3-galactose reduce the humoral barrier to xenotransplantation. Xenotransplantation 20, 27–35 (2013).

61. Laubli, H., et al. Engagement of myelomonocytic Siglecs by tumor-associated ligands modulates the innate immune response to cancer. Proc Natl Acad Sci U S A 111, 14211–14216 (2014).

62. (!!! INVALID CITATION !!! [20]).

63. Carlin, A.F., et al. Molecular mimicry of host sialylated glycans allows a bacterial pathogen to engage neutrophil Siglec-9 and dampen the innate immune response. Blood 113, 3333–3336 (2009).

64. Khatua, B., Bhattacharya, K. & Mandal, C. Sialoglycoproteins adsorbed by Pseudomonas aeruginosa facilitate their survival by impeding neutrophil extracellular trap through siglec- 9. J Leukoc Biol 91, 641–655 (2012).

65. Li, Q. & Lan, P. Activation of immune signals during organ transplantation. Signal Transduct Target Ther 8, 110 (2023).

66. French, B.M., Sendil, S., Pierson, R.N., 3rd & Azimzadeh, A.M. The role of sialic acids in the immune recognition of xenografts. Xenotransplantation 24(2017).

67. Khatua, B., Roy, S. & Mandal, C. Sialic acids siglec interaction: a unique strategy to circumvent innate immune response by pathogens. Indian J Med Res 138, 648–662 (2013).

68. Forgione, R.E., et al. Unveiling Molecular Recognition of Sialoglycans by Human Siglec- 10. iScience 23, 101231 (2020).

69. Kivi, E., et al. Human Siglec-10 can bind to vascular adhesion protein-1 and serves as its substrate. Blood 114, 5385–5392 (2009).

70. Daly, J., Carlsten, M. & O’Dwyer, M. Sugar Free: Novel Immunotherapeutic Approaches Targeting Siglecs and Sialic Acids to Enhance Natural Killer Cell Cytotoxicity Against Cancer. Front Immunol 10, 1047 (2019).

71. Boelaars, K., et al. Unraveling the impact of sialic acids on the immune landscape and immunotherapy efficacy in pancreatic cancer. J Immunother Cancer 11(2023).

72. Scalea, J., Hanecamp, I., Robson, S.C. & Yamada, K. T-cell-mediated immunological barriers to xenotransplantation. Xenotransplantation 19, 23–30 (2012).

73. Goto, M., et al. Dissecting the instant blood-mediated inflammatory reaction in islet xenotransplantation. Xenotransplantation 15, 225–234 (2008).

74. Blaum, B.S., et al. Structural basis for sialic acid-mediated self-recognition by complement factor H. Nat Chem Biol 11, 77–82 (2015).

75. Klaus, C., Liao, H., Allendorf, D.H., Brown, G.C. & Neumann, H. Sialylation acts as a checkpoint for innate immune responses in the central nervous system. Glia 69, 1619–1636 (2021).

76. Friedman, D.J., et al. Cutting Edge: Enhanced Antitumor Immunity in ST8Sia6 Knockout Mice. J Immunol 208, 1845–1850 (2022).

77. Forte, P., Baumann, B.C., Lilienfeld, B., Schneider, M.K.J. & Seebach, J.D. PREVENTION OF NK CELL-MEDIATED CYTOTOXICITY IN PIG-TO-HUMAN XENOTRANSPLANTATION. Transplantation 78, 21 (2004).

78. Lopez, K.J., et al. Strategies to induce natural killer cell tolerance in xenotransplantation. Front Immunol 13, 941880 (2022).

79. Aktas, E., Kucuksezer, U.C., Bilgic, S., Erten, G. & Deniz, G. Relationship between CD107a expression and cytotoxic activity. Cell Immunol 254, 149–154 (2009).

80. del Rio, M.L., Seebach, J.D., Fernandez-Renedo, C. & Rodriguez-Barbosa, J.I. ITIM-dependent negative signaling pathways for the control of cell-mediated xenogeneic immune responses. Xenotransplantation 20, 397–406 (2013).

81. Hamerman, J.A., et al. Cutting edge: inhibition of TLR and FcR responses in macrophages by triggering receptor expressed on myeloid cells (TREM)-2 and DAP12. J Immunol 177, 2051–2055 (2006).

82. Noguchi, Y., et al. Human TIGIT on porcine aortic endothelial cells suppresses xenogeneic macrophage-mediated cytotoxicity. Immunobiology 224, 605–613 (2019).

83. Anand, R.P., et al. Design and testing of a humanized porcine donor for xenotransplantation. Nature 622, 393–401 (2023).

84. Wang, J., et al. Human neural stem cell-derived artificial organelles to improve oxidative phosphorylation. Nat Commun 15, 7855 (2024).

85. Deng, S., Zhang, Y., Shen, S., Li, C. & Qin, C. Immunometabolism of Liver Xenotransplantation and Prospective Solutions. Adv Sci (Weinh) 12, e2407610 (2025).

86. Chen, W., et al. Disulfiram treatment suppresses antibody-producing reactions by inhibiting macrophage activation and B cell pyrimidine metabolism. Commun Biol 7, 488 (2024).

87. Wen, L. & Han, Z. Identification and validation of xenobiotic metabolism-associated prognostic signature based on five genes to evaluate immune microenvironment in colon cancer. J Gastrointest Oncol 12, 2788–2802 (2021).

88. Dobin, A., et al. STAR: ultrafast universal RNA-seq aligner. Bioinformatics 29, 15–21 (2013).

89. Breese, M.R. & Liu, Y. NGSUtils: a software suite for analyzing and manipulating next-generation sequencing datasets. Bioinformatics 29, 494–496 (2013).

90. Liao, Y., Smyth, G.K. & Shi, W. featureCounts: an efficient general purpose program for assigning sequence reads to genomic features. Bioinformatics 30, 923–930 (2014).

91. Robinson, M.D., McCarthy, D.J. & Smyth, G.K. edgeR: a Bioconductor package for differential expression analysis of digital gene expression data. Bioinformatics 26, 139–140 (2010).

92. Mootha, V.K., et al. PGC-1alpha-responsive genes involved in oxidative phosphorylation are coordinately downregulated in human diabetes. Nat Genet 34, 267–273 (2003).

93. Subramanian, A., et al. Gene set enrichment analysis: a knowledge-based approach for interpreting genome-wide expression profiles. Proc Natl Acad Sci U S A 102, 15545–15550 (2005).

